# Transcription-dependent heterochromatin at the *Xist* promoter shapes the random choice of the inactive X chromosome

**DOI:** 10.64898/2026.07.22.737064

**Authors:** Eleni Kanata, Ingrid Pelaez-Conde, Gemma Noviello, Ilona Dunkel, Lena Milanowska, Till Schwämmle, Melissa Bothe, Rutger A.F. Gjaltema, Edda G. Schulz

**Affiliations:** Systems Epigenetics, Otto Warburg Laboratories, Max Planck Institute for Molecular Genetics, 14195 Berlin, Germany; Freie Universität Berlin, Department of Biology, Chemistry, Pharmacy, 14195 Berlin, Germany

## Abstract

In female mammals, Xist, the master regulator of X-chromosome inactivation (XCI), is expressed monoallelically. This pattern is established during early embryonic development, when the active *Xist* allele is chosen at random in each cell. How this choice is made remains incompletely understood. Combining knock-down and overexpression strategies in differentiating mouse embryonic stem cells, which recapitulate the onset of random XCI, we identify a role for the repressive chromatin mark H3K9me3 in the XCI initiation. We show that H3K9me3 accumulates at the promoter-proximal region of the silent *Xist* allele in female cells as monoallelic expression is established. Unexpectedly, this accumulation requires prior transcription of Xist itself, likely during the initial phase of upregulation, when Xist is frequently transcribed in male cells and from both X chromosomes in females. A repressive function of Xist-dependent H3K9me3 accumulation is supported by our finding that premature, transient Xist overexpression primes an allele for future silencing and skews the choice of the inactive X. Xist-dependent H3K9me3 recruitment does not require its antisense transcript Tsix, which can nonetheless enhance subsequent maintenance of the mark. In addition, the X-linked Xist activator RNF12 counteracts H3K9me3 formation, independently of its known target REX1. Our results thus point to facultative heterochromatin formation as a key contributor to choice at the onset of XCI, where activating and repressing mechanisms are intertwined to establish monoallelic Xist expression.

## Introduction

Female mammals silence one of their X chromosomes to ensure dosage compensation for X-linked genes between the sexes (Loda *et al*, 2022). This process of X-chromosome inactivation (XCI) is orchestrated by the long non-coding RNA Xist, which is encoded on the X chromosome. During early embryonic development, monoallelic expression of Xist is established, where the gene is expressed from exactly one X chromosome in each cell. Xist RNA accumulates on the X chromosome *in cis* and recruits repressive factors that lead to the silencing of most X-chromosomal genes (Loda *et al*, 2022). The choice of the *Xist*-expressing allele is random in each cell, resulting in a mosaic pattern, where half of the cells silence the paternal and half the maternal X chromosome. Subsequently, the choice of the *Xist*-expressing allele is stably maintained and inherited through all further cell divisions, attesting to the presence of epigenetic memory (Kanata *et al*, 2024). How female cells ensure monoallelic Xist expression is not yet fully understood.

Random XCI is established at the epiblast stage of mouse development between embryonic day E4.75 and E6.5 (Shiura & Abe, 2019; Cheng *et al*, 2019). While Xist expression is strictly monoallelic at E6.5, it is detected from both alleles in 15-20% of cells at earlier time points (Shiura & Abe, 2019; Mutzel *et al*, 2019; Sousa *et al*, 2018). These dynamics can be recapitulated *in vitro* in differentiating mouse embryonic stem cells (ESCs) from 2i-supplemented media (Sousa *et al*, 2018; Guyochin *et al*, 2014; Mutzel *et al*, 2019; Pacini *et al*, 2021). We have previously shown that differences in chromatin state between the active and silent *Xist* allele are restricted to the promoter-proximal region of Xist (Gjaltema *et al*, 2022). This region contains the main promoter (P1) and a promoter-proximal enhancer located within the first exon, which we named regulatory element 57 (RE57) and which also harbors a secondary promoter P2. This region is enriched for active histone marks on the *Xist*-expressing allele but is covered by the repressive modification H3K9me3 on the *Xist*-silent allele in XO/XY cells upon differentiation. In support of a role of H3K9me3 in Xist regulation, we recently identified the H3K9 methyltransferase SETDB1 as an Xist repressor in a CRISPR screen (Schwämmle *et al*, 2025).

H3K9me3 is mainly associated with constitutive heterochromatin at repeat-rich regions, especially centromeres and telomeres (Saksouk *et al*, 2015), but also with facultative heterochromatin, for example at the promoters of lineage-specific and developmental genes (Wang *et al*, 2018; Burton *et al*, 2020; Nicetto *et al*, 2019). *Xist* thus constitutes an interesting model to study the role of H3K9me3 in gene regulation. H3K9me3 is also a hallmark of other classes of monoallelic genes, such as imprinted genes and olfactory receptors (Kanata *et al*, 2024). Read-write feedback loops enable the mark to spread and to be maintained upon cell division, therefore mediating epigenetic memory (Grewal, 2023). H3K9me3 is closely linked to DNA methylation, since both chromatin marks mutually reinforce each other (Janssen & Lorincz, 2022; Tatarakis *et al*, 2025; Liu *et al*, 2025). H3K9me3 and DNA methylation are thus prime candidates for maintaining the monoallelic expression of Xist. Accordingly, the Xist promoter-proximal region exhibits asymmetric DNA methylation with the *Xist*-expressing allele being hypomethylated in somatic cells (Norris *et al*, 1994; Beard *et al*, 1995; McDonald *et al*, 1998). However, DNA methylation-deficient embryos only show a minimal derepression of Xist (Panning & Jaenisch, 1996; Sado *et al*, 2004), suggesting that H3K9me3 could be compensating.

While various trans-acting Xist regulators have been described, none has been associated with regulation of heterochromatin at the locus. Most notably, the transcription factor (TF) REX1 (ZFP42) acts as an Xist repressor through binding to RE57 (Gontan *et al*, 2012, 2018), where it antagonizes binding of the Xist activator YY1 (Makhlouf *et al*, 2014). REX1 protein levels are controlled by the E3 ubiquitin ligase RNF12 (RLIM), which targets REX1 for degradation (Gontan *et al*, 2012). Recently we identified additional Xist regulators using a CRISPR screen, including several repressors (Schwämmle *et al*, 2025). It remains unknown whether any of them function by promoting heterochromatin formation at *Xist*.

In previous studies, H3K9me3 deposition at the *Xist* gene has been linked to transcription of Xist’s repressive antisense transcript Tsix. A truncation of *Tsix* results in loss of H3K9me3 and DNA methylation at the promoter region in ESCs grown in conventional media, where the marks accumulate without differentiation (Navarro *et al*, 2006). Conversely, ectopic Tsix expression in epiblast stem cells induces accumulation of DNA methylation and, to a lesser extent, H3K9me3 (Ohhata *et al*, 2021). Mechanistically, Tsix transcription recruits H3K36me3, which exerts repression in part through promoting DNA methylation (Ohhata *et al*, 2015). How Tsix promotes H3K9me3 remains unclear, but increased DNA methylation might support maintenance of the mark. Apart from Tsix, also Xist transcription itself has recently been reported to promote H3K9me3 within its own gene body through co-transcriptional recruitment of the human silencing hub (HUSH) complex, which recruits SETDB1 (Almeida *et al*, 2026). A role for Xist transcription in establishing H3K9me3 is difficult to reconcile with the fact that the mark is generally associated with the single, silent *Xist* allele in cells with a single X chromosome (XO/XY). Moreover, whether heterochromatin at *Xist* is required for maintenance of monoallelic Xist expression, or is also involved in the initial choice process remains to be investigated.

Here we set out to characterize the regulation and functional role of heterochromatin formation at the *Xist* locus during the initiation phase of XCI. We find that heterochromatin is formed around the same time when monoallelic Xist expression is established, and is strongly biased to the *Xist*-silent allele. We show that Xist transcription is required for full H3K9me3 recruitment during differentiation, in both XX and XO cells, and that Xist transcription can prime an allele for later repression. This suggests an unexpected role for transient Xist transcription in initiating heterochromatin formation at the *Xist*-silent allele. Tsix is not required for Xist-mediated H3K9me3 deposition but can promote maintenance of the mark. Moreover, we find that RNF12 counteracts H3K9me3 and DNA methylation at the *Xist* promoter region, independently of its characterized target REX1, and that Xist repressors ZFP281 and ZFP36L1 promote heterochromatin accumulation at the locus. Our findings, for the first time, implicate heterochromatin formation at *Xist* in the choice process that initiates XCI.

## Results

### Timing of DNA methylation and H3K9me3 at the *Xist* promoter coincides with establishment of monoallelic expression

We have previously shown that H3K9me3 accumulates around the *Xist* promoter at the onset of XCI on the *Xist*-silent allele in mouse XO and XY cells, but not on the *Xist*-expressing allele in XX cells (Gjaltema *et al*, 2022) (Fig. 1a-b, Suppl. Fig. 1a). The mark is also present *in vivo* at the onset of random XCI, as ChIP-seq data from sex-mixed mouse embryos (Wang *et al*, 2018) revealed an H3K9me3 domain around the *Xist* promoter between E3.5 and E7.5 (Suppl. Fig. 1b). It has not yet been tested whether DNA methylation, which is often associated with H3K9me3, exhibits a similar timing and pattern as H3K9me3. We therefore set out to characterize the dynamics of DNA methylation using whole-genome bisulfite sequencing (WGBS) in the same experimental system, and compare them to our previously generated H3K9me3 data (Gjaltema *et al*, 2022). We profiled the *Xist*-silent allele using XO ESCs, and the *Xist*-expressing allele using XX_ΔXIC-B6_ ESCs (Mutzel *et al*, 2025) (Fig. 1b), both generated from an ESC line (TX1072) derived from a cross between two distantly related mouse strains (B6/Cast). XX_ΔXIC-B6_ ESCs carry a 773kb deletion surrounding *Xist* on the B6 allele, leaving only the *Xist*-expressing Cast allele. We analyzed naive ESCs (2i/FBS/LIF) and cells differentiated by 2i/LIF withdrawal for 2 and 4 days. In XX cells, Xist upregulation occurs between differentiation day 1 and 2, transient biallelic expression is resolved by day 3 and X-linked gene silencing is completed by day 4 (Pacini *et al*, 2021). In XO cells, Xist is also transiently expressed at low levels between day 1 and 2 of differentiation (Suppl. Fig. 1c), similar to XY cells (Sousa *et al*, 2018).

**Figure 1:**
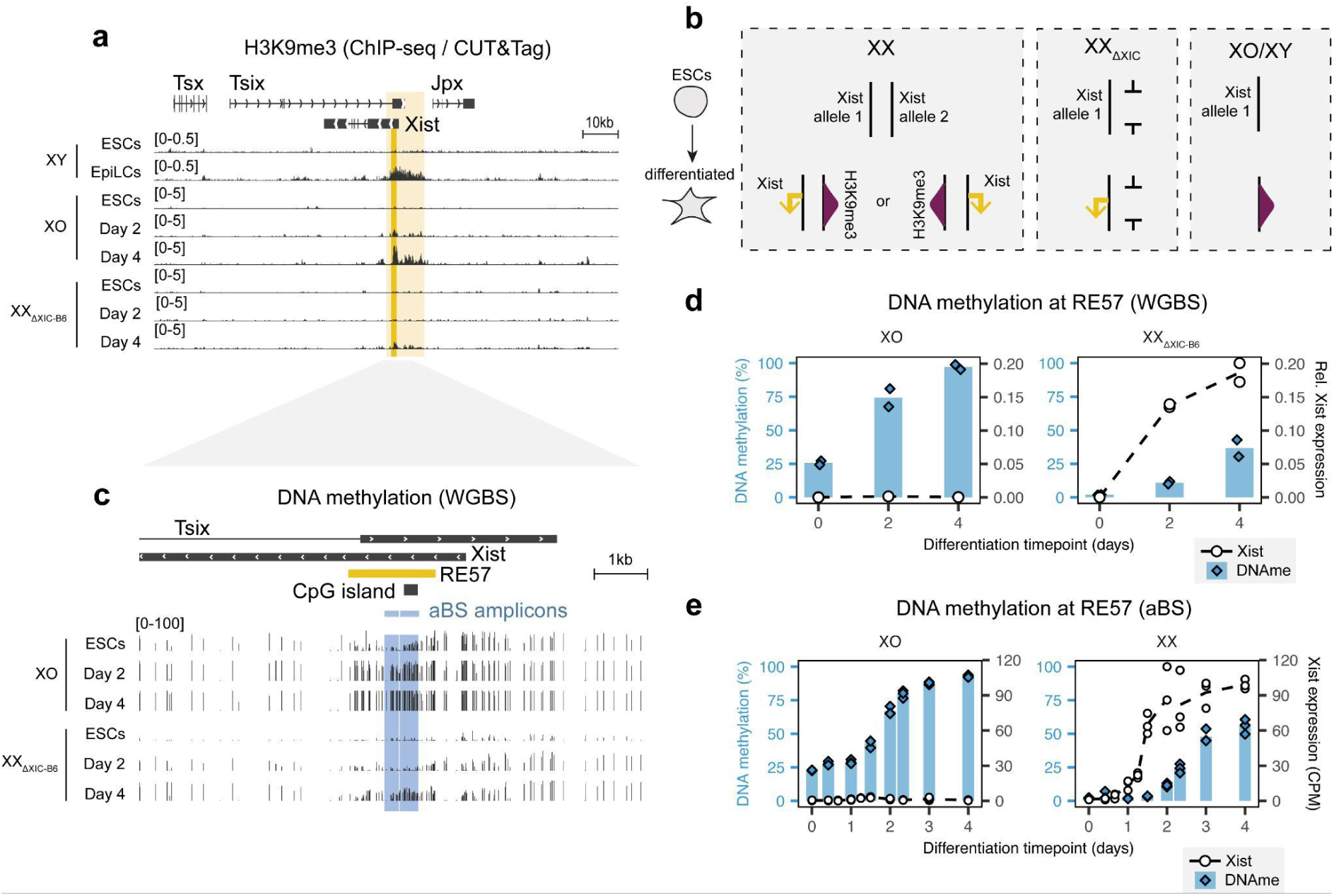
Dynamics of heterochromatin marks at the *Xist* promoter-proximal region. (a) H3K9me3 XY ChIP-seq tracks from (Bleckwehl *et al*, 2021). H3K9me3 CUT&Tag in differentiating XX_ΔXIC-B6_ and XO cells (Gjaltema *et al*, 2022), bigwig tracks, average of 2 biological replicates. Light yellow box: H3K9me3-marked area around the *Xist* promoter. Dark yellow box: RE57. Region shown: chrX:103,407,345-103,550,559 (mm10). (b) Schematic illustration of cell lines used in this study. (c) DNA methylation levels at individual CpGs as the percentage of methylated reads, measured by WGBS. Two biological replicates averaged per condition. XX_ΔXIC-B6_ from (Mutzel *et al*, 2025), XO data from this study. Amplicons used in amplicon bisulfite sequencing (aBS) marked in blue. Region shown: chrX:103,477,061-103,486,581 (mm10). (d) DNA methylation in RE57 for the data shown in c. Mean (bars) of two biological replicates (blue diamonds). Xist expression (RT-qPCR) from the same experiment is shown (white marks) (Gjaltema *et al*, 2022). (e) DNA methylation levels measured by aBS. The mean of the two amplicons is plotted (blue bars) for three biological replicates (blue diamonds). Xist expression as measured by RNA-sequencing (CPM) from the same experiment (Schwämmle *et al*, 2025) is overlaid (white marks).

The WGBS data revealed that the region surrounding the CpG island within *Xist’s* promoter-proximal enhancer RE57 exhibited low DNA methylation at the naive stage in both cell lines (Fig. 1c-d). During differentiation, RE57 remained hypomethylated in XX_ΔXIC-B6_ cells, while it gained high methylation levels in XO cells (Fig. 1c-d). To determine the timing of DNA methylation in relation to Xist expression, we performed amplicon bisulfite sequencing (aBS) during a differentiation time course with higher time resolution. The designed amplicons cover the CpG island and the neighboring YY1 binding sites, which are occupied by YY1 only when unmethylated (Makhlouf *et al*, 2014). WGBS and aBS results were highly concordant (Suppl. Fig. 1d). We compared XO to XX ESCs (TX1072), which upregulate Xist from one randomly chosen X chromosome and are thus expected to reach around 50% of DNA methylation. DNA methylation increased in a switch-like manner in both cell lines, starting around day 1 in XO cells and around day 2 in XX cells (Fig. 1e, Suppl. Fig. 1e). Heterochromatin formation thus occurs after initial Xist upregulation (day 1), when monoallelic expression is established and any spurious Xist expression (biallelic in XX, expression in XO) is silenced (day 3) (Pacini *et al*, 2021).

### H3K9me3 specifically marks the *Xist*-silent allele in XX cells after XCI establishment

Since we profiled the *Xist*-silent allele only in XO cells, we next set out to assess whether the *Xist*-silent allele in XX cells would exhibit a similar chromatin state. We developed a new approach to probe the chromatin landscape of the two *Xist* alleles in XX cells separately without interfering with the random choice of XCI. Through tagging each allele of an X-linked gene with a different fluorescent protein, we could sort cells based on which X chromosome they silenced as a proxy for the active *Xist* allele, and perform allele-specific H3K9me3 profiling. Within our XX ESC line (TX1072), we tagged the *Maged1* gene with mScarlet-I3 and mStayGold on the B6 and Cast allele, respectively (Fig. 2a, Suppl. Fig. 2a-d). Xist induction dynamics in the tagged cell line (TX-Maged1-2tag) were comparable to the parental line, although Xist RNA levels were slightly reduced and some variation in differentiation dynamics were observed (Suppl. Fig. 2e-f). Random XCI results in silencing of Maged1 on one allele, and therefore in a reduction of either mStayGold or mScarlet-I3 (Fig. 2b). After differentiation for 4 days, we sorted three populations by fluorescence-activated cell sorting (FACS): cells that had silenced the B6 allele (Xist-B6), those that had silenced the Cast allele (Xist-Cast), and cells that still carried two active X chromosomes (Xist-Off) (Fig. 2b, Suppl. Fig. 2g). RNA Fluorescence In Situ Hybridization (RNA-FISH) for Xist confirmed high purity of the sorted populations, since Xist-B6 and Xist-Cast populations contained between 92% and 97% cells with monoallelic Xist expression, while 98% of Xist-Off cells did not express Xist (Fig. 2c).

**Figure 2:**
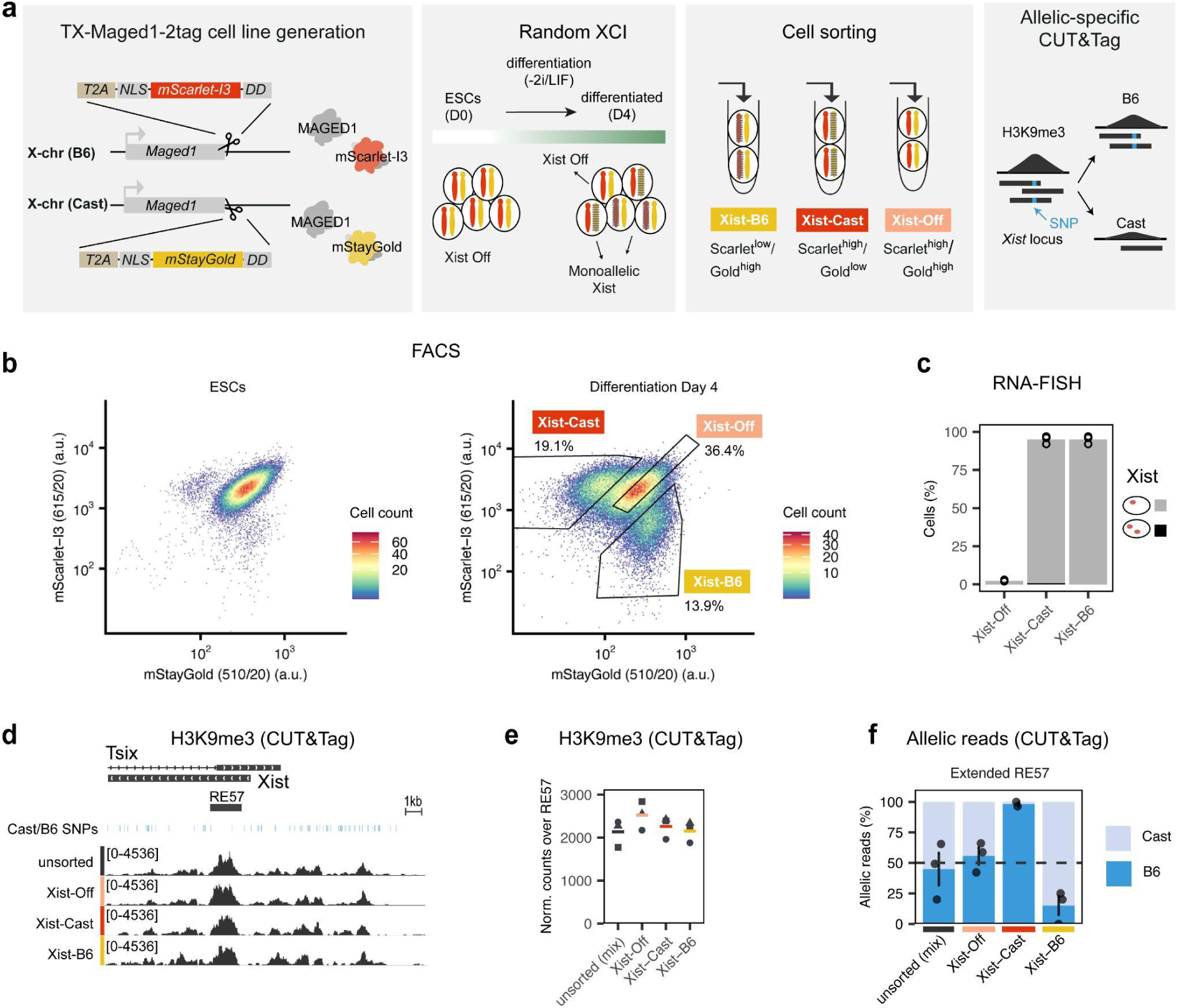
Monoallelic H3K9me3 deposition in differentiating XX ESCs at the *Xist* promoter-proximal region. (a) Scheme of TX-Maged1-2tag cell line and experimental setup. X-linked gene *Maged1* is C-terminally tagged by insertion of a fluorescent protein. T2A: self-cleaving peptide, NLS: nuclear localization signal, DD: destabilizing domain FKBP12^F36V-L106P^ (FK506-Binding Protein 12, carrying mutations F36V and L106P). Upon differentiation, cells are sorted into the indicated populations, followed by allele-specific CUT&Tag. (b) Flow cytometry analysis of TX-Maged1-2tag cells at the ESC stage (left) and after 4 days of differentiation (right). Hexagonal heatmap of cell counts in 256 bins. The sorting gates are indicated together with the percentage of cells in each gate. (c) RNA-FISH for Xist in the cell populations sorted with the gates shown in (b). 100 cells were counted per sample. The mean of n=3 independent replicates (dots) is shown. (d) H3K9me3 CUT&Tag tracks from each sorted population (mean of 3 biological replicates for each condition, normalized bigwig). Reads from both alleles in the range chrX:103,475,052-103,493,155 (mm10). (e) Normalized H3K9me3 CUT&Tag reads in RE57. Symbols represent individual biological replicates and horizontal lines the mean. (f) Allelic fraction of SNP-covering CUT&Tag reads spanning extended RE57 region (chrX:103,479,041-103,482,895). Dark blue (bottom) B6 reads, light blue (top) Cast reads. Dots show individual replicates and vertical lines standard deviation.

We performed CUT&Tag for H3K9me3 for unsorted cells and for all three sorted populations. We then quantified the overall signal in RE57 (Fig. 2d, e) as well as the contribution of the two alleles at a 3.9 kb SNP-rich region around RE57 (Fig. 2f). While both alleles contributed equally to the signal in unsorted and Xist-Off cells, the vast majority of reads mapped to the B6 allele in Xist-Cast cells and to the Cast allele in Xist-B6 cells, supporting the specific deposition of the mark on the silent *Xist* allele. Notably, the overall H3K9me3 signal appeared to be increased in Xist-Off cells compared to the other populations, suggesting biallelic H3K9me3 marking (Fig. 2e). These results show that H3K9me3 indeed accumulates on the *Xist*-silent allele in XX cells when XCI is established, and might accumulate on two alleles in cells that fail to stably upregulate Xist.

### Initial Xist transcription primes its future silencing

It has recently been reported that Xist transcription can drive deposition of H3K9me3 when overexpressed with a Doxycycline (Dox)-inducible system or through *Tsix* deletion (Almeida *et al*, 2026). This finding seems contradictory to our observation that H3K9me3 is primarily found at the *Xist*-silent allele. However, since Xist is transiently expressed from both X chromosomes, at least in a percentage of cells in females and from the single X in males during early differentiation (Sousa *et al*, 2018; Mutzel *et al*, 2019; Shiura & Abe, 2019), we hypothesized that initial Xist transcription at the future *Xist*-silent allele, might help to initiate heterochromatin formation.

To test whether endogenous Xist transcription has a role in recruiting H3K9me3 during differentiation, we knocked-down Xist and quantified H3K9me3 and DNA methylation. We used CasTuner, an inducible CRISPRi system we have recently developed, consisting of a catalytically dead Cas9 (dCas9) fused to the repressor domain HDAC4, that encodes a histone deacetylase, and to a dTAG-controlled conditional degron (Noviello *et al*, 2023) (Fig. 3a, Suppl. Fig. 3a). TX1072 ESCs stably expressing CasTuner are maintained in dTAG-containing media, which degrades the system, and knock-down (KD) is induced by dTAG withdrawal (-dTAG). We knocked down Xist in both XX and XO cells, and measured H3K9me3 (C&T) and DNA methylation (aBS) at day 4 of differentiation.

**Figure 3:**
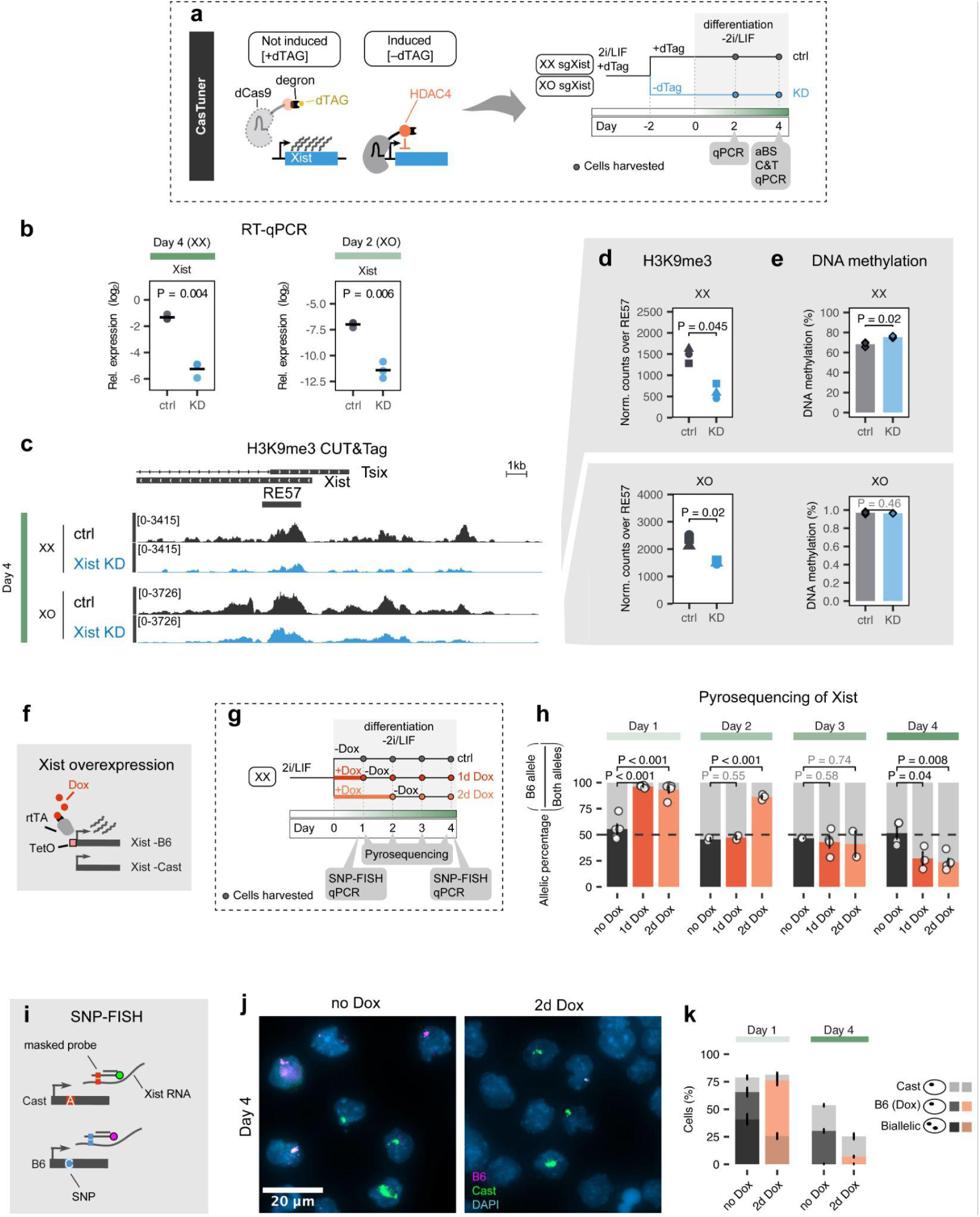
Transient Xist transcription primes locus for later repression. (a) Experimental setup for Xist KD in XX and XO cells with CasTuner. (b) RT-qPCR data for Xist expression. Horizontal lines indicate the mean of 3 biological replicates (dots). ctrl: +dTAG, KD: -dTAG. Two-sided unpaired T-test, p-value black if <0.5. (c) H3K9me3 CUT&Tag tracks (mean of 3 biological replicates for each condition, normalized bigwig). (d) Number of normalized reads covering RE57. Each of the three shapes corresponds to a biological replicate. Two-sided paired T-test. (e) DNA methylation measured by aBS. The mean of the two amplicons is plotted for three biological replicates (diamonds). Two-sided paired T-test. (f) Illustration of Xist overexpression with Dox in the XX ES cell line. The B6 allele is Dox-inducible. Dox: doxycycline, TetO: Tetracycline-operator that renders the *Xist* promoter Dox inducible, rtTA: reverse tetracycline-controlled trans-activator. (g) Timeline for transient Xist overexpression. Dox is washed out either after 1 or 2 days of treatment. (h) Pyrosequencing of Xist RNA. Percentage of reads coming from the B6 (overexpressed) allele at each differentiation day and treatment are plotted. Error bar shows standard deviation. Two-sided unpaired T-test. Between 2 and 5 replicates for each condition. (i) Illustration of SNP-FISH. In this approach 28 RNA-FISH probes are used, each containing a SNP. To allow specific binding to one sequence variant, part of the probe is initially masked through a complementary shorter oligo. (j) SNP-FISH microscopy images at day 4 of differentiation with no or 2-day Dox treatment. (k) Allelic Xist signal from SNP-FISH of three biological replicates at day 1 and 4 of differentiation, with 2 days of Dox treatment (orange) or without treatment (gray). Error bar shows standard deviation. Dark colors correspond to biallelic cells, lighter colors to monoallelic.

We assessed KD efficiency at day 4 for XX cells and at day 2 for XO cells, when Xist expression is maximal (Suppl. Fig. 1c), revealing a reduction of Xist RNA levels by 93-95% (Fig. 3b). Unexpectedly, we also observed a ∼2-fold reduction in Tsix levels in both cell lines (Suppl. Fig. 3b). This suggests that targeting CasTuner to the *Xist* promoter, which lies within a Tsix exon, might also interfere with Tsix elongation. Upon Xist KD, the levels of H3K9me3 at the locus were significantly reduced by 58% in XX and by 35% in XO cells compared to the control (+dTAG) (Fig. 3c-d), supporting a role of Xist transcription in H3K9me3 deposition at the *Xist*-silent allele. A decrease in DNA methylation was not observed in either cell line, and in XX cells we even observed a slight increase (Fig. 3e). These results suggest that the future *Xist*-silent allele is transiently transcribed in a substantial fraction of cells and that this has an unexpected functional role even in cells with a single X chromosome.

If Xist transcription can indeed initiate heterochromatin formation and thereby prime the locus to become the silent *Xist* allele, we reasoned that premature Xist transcription on one allele would modulate the choice of the inactive X. To test this prediction, we made use of the Dox-inducible promoter that is present upstream of the *Xist* transcription start site (TSS) on the B6 allele in the TX1072 ESCs to transiently induce Xist transcription (Fig. 3f). We treated cells with Dox only for the first one or two days of differentiation and analyzed Xist expression after Dox washout up to day 4 (Fig. 3g, Suppl. Fig. 3c). For an allele-resolved analysis, we performed pyrosequencing of a SNP within the Xist RNA to quantify the contribution of each allele to the amount of Xist RNA produced by the cell population. While the B6 allele contributed ∼50% of Xist RNA at all timepoints in the untreated control, Dox treatment increased that fraction to 80-90% (Fig. 3h). One day after Dox washout, the B6 contribution was reduced to 50%, comparable to the untreated control. Surprisingly, it was further reduced to 24-27% at day 4 of differentiation, thus resulting in inverse skewing of Xist expression compared to acute Dox treatment (day 1). The fact that the allele where Xist was prematurely induced produces a minority of Xist RNA at a later timepoint suggests that Xist transcription indeed primes the allele for later repression, whether that is complete silencing or lower transcription levels.

That fact that only 24% of the total Xist RNA in the population is transcribed from the B6 chromosome, might suggest that the B6 allele is indeed silent in the majority of the cells. However, an alternative explanation could be that the Dox pulse results in continued expression, but at reduced levels once Dox is washed out. To distinguish these possibilities, we established SNP FISH for Xist for allele-specific detection of Xist RNA on the single-cell level (Levesque 2013). We designed probes for both the B6 and the Cast allele and used them to assign each Xist signal to one or the other allele (Fig. 3i-k). This revealed that in Dox-treated cells at day 1, 50% of cells expressed the B6 allele only, 5% the Cast allele only, and 26% of cells expressed both alleles (Fig. 3k). At day 4 by contrast, expression was nearly exclusively monoallelic. While the percentage of Cast-Xist cells was comparable in two-day-treated cells and in untreated cells (18% vs 23%), the amount of B6-Xist cells was strongly reduced (7% vs 30%). This shows that transient Dox treatment indeed results in silencing of Xist on the induced allele. As a consequence, the overall amount of Xist-positive cells as well as RNA levels are reduced upon transient Dox treatment (Fig. 3k, Suppl. Fig. 3c). To assess the consequences on X-linked gene silencing, we also measured allelic expression of two X-linked genes, Rnf12 and Fam122b, by pyrosequencing (Suppl Fig. 3d).

Although no major differences were observed, at day 4 the one-day-treated cells exhibited a significantly higher B6 contribution for both genes compared to untreated cells. This shows that a higher percentage of cells have silenced the Cast chromosome in the treated group at day 4, and is in agreement with the SNP-FISH results, where more cells express Xist from the Cast allele. Taken together, transient premature Xist induction skews allelic choice to the opposite allele. This finding is consistent with a model where transient Xist transcription during early differentiation promotes repression of that allele through H3K9me3 recruitment. Our results thus suggest a role for Xist-dependent heterochromatin formation in the choice process.

### Tsix transcription is not required for Xist-dependent H3K9me3 accumulation

While our data suggest that initial Xist expression can prime the locus for silencing, it is not clear how H3K9me3 is later restricted to the silent *Xist* allele. A major difference between the alleles is the activity of Tsix, which is silenced along with the rest of the X chromosome on the *Xist*-expressing allele (Fig. 4a). We hypothesized that Xist expression might only be able to instruct H3K9me3 deposition when co-transcribed with Tsix. This would, for example, allow the formation of double-stranded RNA, which in some organisms can trigger RNAi-mediated heterochromatin formation (Reyes-Turcu & Grewal, 2012). To test the necessity of Tsix in Xist-directed heterochromatin, we ectopically induced Xist transcription with Dox, in naive cells, where it is normally silent, both in presence and absence of Tsix transcription. To knock down Tsix, we co-targeted two of its promoters with CasTuner (Fig. 4a-b). We induced Xist overexpression in the naive state for two days in the presence and absence of dTAG to induce Tsix KD (Fig. 4c,d). We observed a strong recruitment of H3K9me3 at RE57 and the surrounding locus, especially downstream of the *Xist* TSS (Fig. 4e-f). This H3K9me3 accumulation was unaffected by a 68% reduction of Tsix transcription (Fig. 4d-f). Instead, Tsix KD appeared to rather increase H3K9me3 levels, potentially due to increased Xist transcription. Moreover, Xist-mediated H3K9me3 recruitment occurred only *in cis*, since it was restricted to the B6 allele based on allele-specific CUT&Tag analysis, also arguing against an RNAi-dependent mechanism (Suppl. Fig. 4a). These results show that co-transcription of Xist and Tsix is not required for H3K9me3 deposition and that Xist transcription alone can recruit high levels to the locus *in cis*, independently of differentiation.

**Figure 4:**
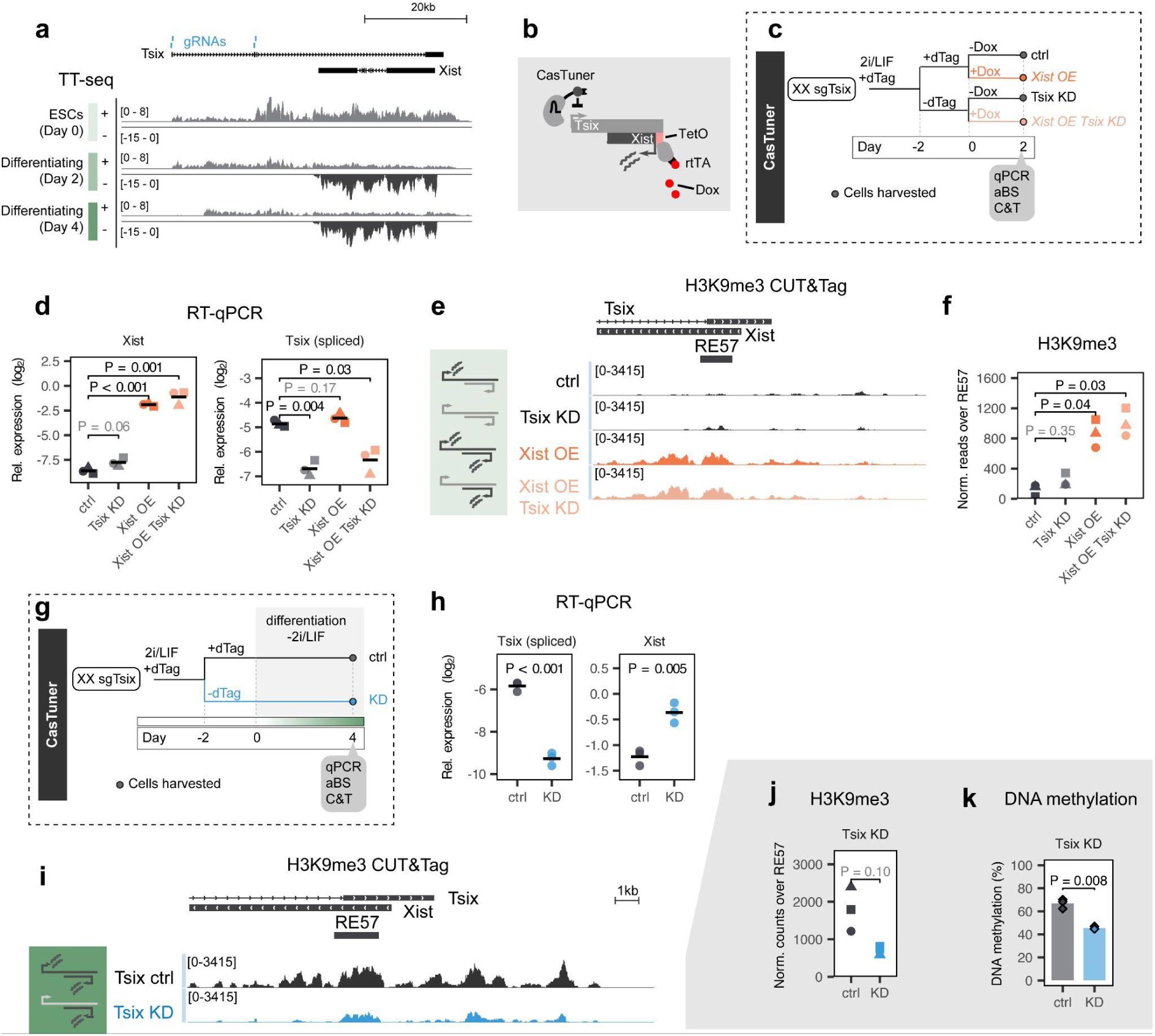
Role of Tsix in heterochromatin formation at *Xist*. (a) Nascent transcription (TT-seq) data of XX_ΔXIC-B6_ from (Gjaltema *et al*, 2022), average of 2 biological replicates. In blue the gRNAs that target the *Tsix* TSSs in the Tsix KD. Region plotted: chrX:103,421,685-103,490,000 (mm10). (b) Illustration of simultaneous Tsix KD and Xist overexpression in the Tsix CasTuner cell line (sgTsix). Dox: doxycycline, TetO: Tetracycline-operator that renders the *Xist* promoter Dox inducible, rtTA: reverse tetracycline-controlled transactivator. (c) Experimental timeline used in d-f. OE: Overexpression. (d) RT-qPCR data for Xist and Tsix expression. ctrl: +dTAG, KD: -dTAG. Two-sided unpaired T-test. Horizontal lines represent the mean of three biological replicates (dots). (e) H3K9me3 CUT&Tag tracks (mean of 3 biological replicates for each condition, normalized bigwig). (f) Normalized reads covering RE57. Each of the three shapes corresponds to a biological replicate. Two-sided paired T-test. (g) Experimental scheme for testing the role of Tsix transcription in H3K9me3 recruitment used in h-k. (h) RT-qPCR data for Xist and Tsix expression. ctrl: +dTAG, KD: -dTAG. Two-sided unpaired T-test. Horizontal lines represent the mean of three biological replicates (dots). (i) H3K9me3 CUT&Tag tracks (mean of 3 biological replicates for each condition, normalized bigwig). (j) Normalized reads covering RE57. Each of the three shapes corresponds to a biological replicate. Two-sided paired T-test. (k) DNA methylation measured by aBS. The mean of the two amplicons is plotted for three biological replicates (diamonds). Two-sided paired T-test.

Although Tsix transcription is not required for H3K9me3 nucleation, it might instead contribute to its maintenance, for example through promoting DNA methylation. In this case, Tsix would enrich the mark on alleles with low or no Xist transcription, where it is preferentially transcribed. To test a role of Tsix in heterochromatin formation, we performed a Tsix KD during differentiation and measured H3K9me3 and DNA methylation levels (Fig. 4g). This approach resulted in a 90% KD and a 1.8-fold increase of Xist expression at day 4 (Fig. 4h). The levels of H3K9me3 at RE57 were reduced by 60%, although the change was not statistically significant, and it was accompanied by a 30 % reduction in DNA methylation (Fig. 4i-k). These results show that *Tsix* is likely required for proper H3K9me3 and DNA methylation deposition at the *Xist* locus in differentiating ESCs. Tsix transcription, which becomes progressively restricted to the *Xist*-silent allele during differentiation, might thus contribute to the preferential maintenance of H3K9me on that allele.

Given that the H3K9me3-marked *Xist* allele also gains DNA methylation (Fig. 1), we evaluated whether *Xist* overexpression can lead to DNA methylation recruitment in addition to H3K9me3. Here we used an XX_ΔXIC-Cast_ (clone A6) cell line, where the *Xist* locus is deleted on the Cast allele, to specifically observe effects *in cis* on the Dox-induced B6 allele. We induced *Xist* overexpression with Dox for two days in the naive state and during differentiation and measured H3K9me3 with CUT&Tag and DNA methylation with amplicon bisulfite sequencing (Suppl. Fig. 4b-c). In both cases we observed a strong increase of H3K9me3 at the locus (Suppl. Fig. 4d-e). Interestingly, this was not accompanied by DNA methylation in naive cells, but led to increased DNA methylation at day 2 of differentiation (Suppl Fig. 4f). DNA methylation is thus not required for Xist-mediated H3K9me3 recruitment.

In summary, we have found that Xist can induce H3K9me3 accumulation independently of Tsix and independently of DNA methylation. Nevertheless, during differentiation, Tsix is required for full H3K9me3 deposition. Xist and Tsix thus appear to cooperate in promoting H3K9me3 deposition during differentiation.

### Xist activator RNF12 counteracts heterochromatin formation at *Xist*

We have identified an unexpected role for Xist-induced H3K9me3 during the initiation of XCI, and found that its manipulation can skew choice of the inactive X. Previous work has identified an important role for X-linked Xist trans-activators in establishing monoallelic expression and in the choice of the inactive X (Mutzel & Schulz, 2020). We therefore asked whether their function would be in part mediated by manipulating heterochromatin formation at the locus. We therefore investigated whether KD of the potent X-linked Xist activators RNF12 (Jonkers *et al*, 2009) and ZIC3 (Schwämmle *et al*, 2025) would affect heterochromatin formation at RE57 (Fig. 5a). Moreover, we also tested the Xist repressor REX1, which is a known target of RNF12, as well as a set of putative Xist repressors we have recently identified in a CRISPR screen (ZFP280C, ZFP36L1, ZFP281) (Schwämmle *et al*, 2025) (Suppl. Fig. 5).

**Figure 5:**
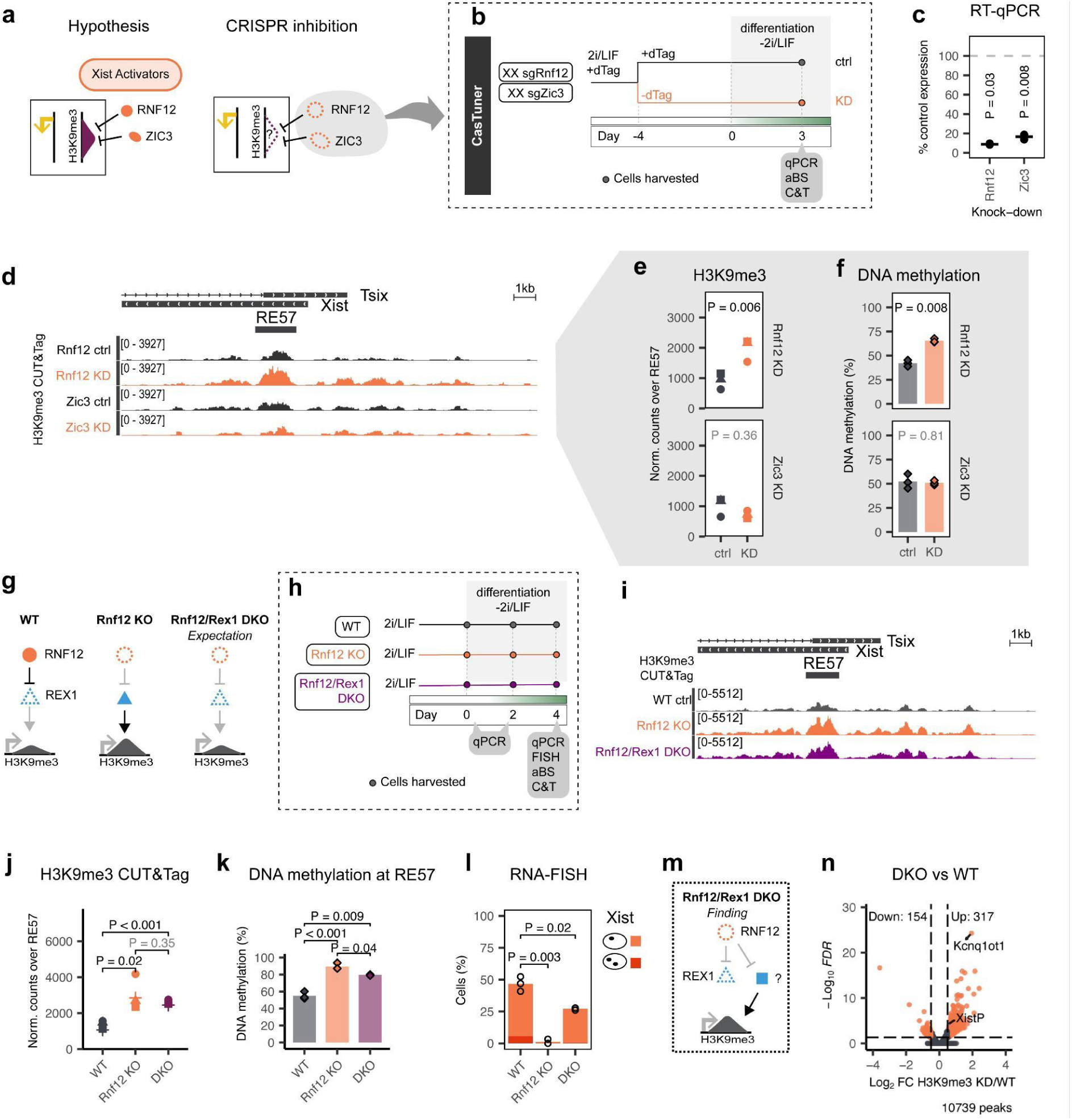
RNF12 protects the *Xist* locus from heterochromatin independently of REX1. (a) Scheme of hypothesis and experiment for activator KD. Xist activators might function by inhibiting H3K9me3 (purple) at the *Xist* locus. This was tested by depleting activators by CRISPRi. (b) Experimental setup for the KD used in c-f. (c) Knock-down efficiency measured by RT-qPCR, as the percentage of target-gene expression in the KD sample relative to the control. The black line represents the mean of the 3 biological replicates. Two-sided unpaired T-test of relative expression of KD vs control. (d) H3K9me3 CUT&Tag tracks (average of 3 biological replicates for each condition, normalized bigwig). (e) Normalized reads covering RE57. Two-sided paired T-test. (f) DNA methylation measured by aBS. The mean of the two amplicons is plotted for three biological replicates (diamonds). Two-sided paired T-test. (g) Hypothesis for REX1-dependent function of RNF12 at H3K9me3 at the *Xist* locus tested in h-n. (h) Experimental design used in i-n. (i) H3K9me3 CUT&Tag tracks (average of 4 biological replicates for WT and Rnf12 KO, 3 for Rnf12/Rex1 DKO, normalized bigwig chrX:103,475,052-103,493,155 mm10). (j) Normalized reads covering RE57. Two-sided unpaired T-test. (k) DNA methylation measured by aBS. The mean of the two amplicons is plotted for three biological replicates (diamonds). Two-sided unpaired T-test. (l) Percentage of Xist-positive cells in RNA-FISH. Monoallelic percentage in orange, biallelic in red. Three biological replicates per condition, 98-151 cells counted per sample. (m) Schematic model of REX1-independent function of RNF12 at H3K9me3 at *Xist*. (n) CUT&Tag volcano plot after differential analysis (diffbind (DESeq2) on epic2-identified H3K9me3 peaks). Orange dots represent peaks that had a significant change, defined as an FDR < 0.05 and an absolute log2 fold change > 0.5. XistP: *Xist* promoter-proximal region

To assess the role of these factors in heterochromatin formation, we depleted them in differentiating XX ESCs through CRISPRi (CasTuner or a dCas9-KRAB system) and quantified H3K9me3 and DNA methylation at RE57 (Fig. 5b, Suppl. Fig. 5b). We confirmed the KD efficiency by RT-qPCR (Fig. 5c, Suppl. Fig. 5c) and verified that the KD did not exhibit major effects on cell differentiation (Suppl. Fig. 5f,h). While the KD of activators generally led to a decrease in Xist RNA, and the repressor KD to an increase, the effects were variable and not always statistically significant (Suppl. Fig. 5d-e,g). Analysis of DNA oligo spike-in controls revealed no genome-wide effect on H3K9me3 levels (Suppl. Fig. 6a-b).

While Zic3 KD had no detectable effect on heterochromatin formation, Rnf12 KD resulted in 2.2-fold increased levels of H3K9me3 compared to the uninduced control (Fig. 5d, e). Also, the levels of DNA methylation at RE57 were significantly increased upon Rnf12 KD over the whole extent of covered CpGs (Fig. 5f, Suppl. Fig. 6c). Surprisingly, we did not observe a clear effect upon Rex1 KD, although this factor is thought to mediate RNF12-dependent Xist regulation (Suppl. Fig. 6d-f). Among the other tested factors, Zfp281 and Zfp36l1 KD resulted in reduced H3K9me3 and DNA methylation at RE57 (Suppl. Fig. 6d-f), suggesting that these factors promote heterochromatin formation. In summary, we have identified a new role for RNF12 in counteracting heterochromatin formation at RE57 and identified two factors, ZFP281 and ZFP36L1, that promote the repressive chromatin state.

### RNF12 regulates heterochromatin independently of REX1

RNF12 is an E3 ubiquitin ligase that regulates *Xist* through the ubiquitination of REX1, resulting in its proteasomal degradation (Gontan *et al*, 2012). The fact that we observed an effect on heterochromatin upon Rnf12, but not upon Rex1 KD raises the question whether this function of RNF12 is mediated by REX1 or not. To answer this question, we used a previously generated set of female ESC lines, either carrying a homozygous deletion of Rnf12 (RNF12-/-, in short Rnf12 KO), or a double deletion of Rnf12 and Rex1 (RNF12-/-REX1-/-, in short DKO) (Gontan *et al*, 2018). If increased REX1 protein levels in Rnf12 KO would mediate H3K9me3 deposition in the absence of RNF12, the additional deletion of REX1 would prevent the increase of H3K9me3 at *Xist* (Fig. 5g).

We differentiated the cells for 4 days and performed CUT&Tag and aBS (Fig. 5h, Suppl. Fig. 7a). Similarly to the Rnf12 KD, the Rnf12 KO gained 2.3-fold higher levels of H3K9me3 at RE57 than the wildtype (WT) control (Fig. 5i-j). The DKO also showed 2 times higher levels of H3K9me3, supporting a REX1-independent role of RNF12 at RE57. Regarding DNA methylation, we observed that levels were increased in both mutant cell lines compared to the WT control, with the increase being more pronounced in the Rnf12 KO (Fig. 5k). To test whether this REX1-independent function of RNF12 contributes to Xist regulation, we analyzed Xist expression in the three cell lines by RNA-FISH and RT-qPCR (Fig. 5l, Suppl. Fig. 7b). Xist upregulation was essentially abolished in RNF12 KO cells, and was only partially rescued in the DKO cell line, where Xist was expressed in 27% of cells compared to 47% in WT cells (Fig. 5l). These observations are in agreement with the original description of the cell lines, which used a different cell culture and differentiation protocol (Gontan *et al*, 2018). The partial rescue of Xist expression in the DKO had been attributed to a slower differentiation of the DKO cell line compared to the WT. However, in our differentiation system, we observed decreased levels of Nanog at day 2 (Suppl. Fig. 7c), rather indicating accelerated differentiation. Therefore, we propose that the lower frequency of Xist-expressing cells in DKO cells arises from increased H3K9me3 deposition. This highlights that RNF12 counteracts heterochromatin at the *Xist* locus in a REX1-independent manner that is relevant for proper Xist upregulation (Fig. 5m).

To investigate whether this REX1-independent function of RNF12 affects other regions besides the *Xist* promoter, we performed a genome-wide analysis of H3K9me3 in the mutant cell lines (Suppl. Table 1). While a comparable number of peaks were lost and gained in the Rnf12 KO, the DKO mainly showed a gain of H3K9me3, as observed at RE57 (Fig. 5n, Suppl. Fig. 7d). The gained peaks in both mutants were highly concordant (Suppl. Fig. 7e). Interestingly, the DKO did not lead to the formation of new peaks but to an increase of H3K9me3 in existing peaks (Suppl. Fig 7f). This suggests that the newly-identified REX1-independent role of RNF12 is affecting the maintenance mechanism rather than the nucleation of heterochromatin. Notably, one of the top H3K9me3-gaining peaks is the long non-coding RNA Kcnq1ot1, which, similarly to Xist, leads to silencing of the surrounding genomic locus *in cis* (Suppl Fig. 7g). This gene is also a target of the HUSH complex (Garland *et al*, 2022), and its elongation is required for its silencing function (Mancini-Dinardo *et al*, 2006). The *Kcnq1ot1* H3K9me3 peak is also affected by the KD of Zfp281 and Zfp36l1, similarly to the *Xist* peak (Suppl Fig. 7g, Suppl. Table 1).

In conclusion, RNF12 counteracts heterochromatin formation at multiple loci, including *Xist*, independently of REX1. The mechanism of the REX1-independent function remains an open question and might involve the ubiquitination of another locus-specific heterochromatin regulator.

## Discussion

Here we describe a new role for H3K9me3 in the initiation of X inactivation. Together with DNA methylation, the mark is recruited to the *Xist* promoter-proximal region around the time when monoallelic expression is established. Using a reporter cell line that allows allele-specific profiling during random XCI without skewing the choice, we show that promoter-proximal heterochromatin accumulates specifically at the *Xist*-silent allele in female cells after establishment of X inactivation. Surprisingly, we found evidence that full H3K9me3 recruitment requires prior transcription of Xist. This suggests that transient Xist expression from the future *Xist*-silent allele promotes initiation of heterochromatin formation *in cis*. While Xist-dependent H3K9me3 recruitment does not require Tsix, the antisense transcript is required for full maintenance of the mark during differentiation. H3K9me3 deposition is in addition promoted by absence of the X-linked Xist activator RNF12 through a mechanism independent of its target REX1. Our results thus suggest a functional role for transient Xist upregulation from all X chromosomes (one in males, two in females) that has previously been described to occur during XCI onset (Shiura & Abe, 2019; Mutzel *et al*, 2019; Sousa *et al*, 2018). We propose that heterochromatin formation is initiated at this stage through Xist-mediated H3K9me3 deposition and through silencing of all *Rnf12* gene copies in a cell. Maintenance of H3K9me3 on the silent *Xist* allele is subsequently promoted by Tsix (Fig. 6).

**Figure 6:**
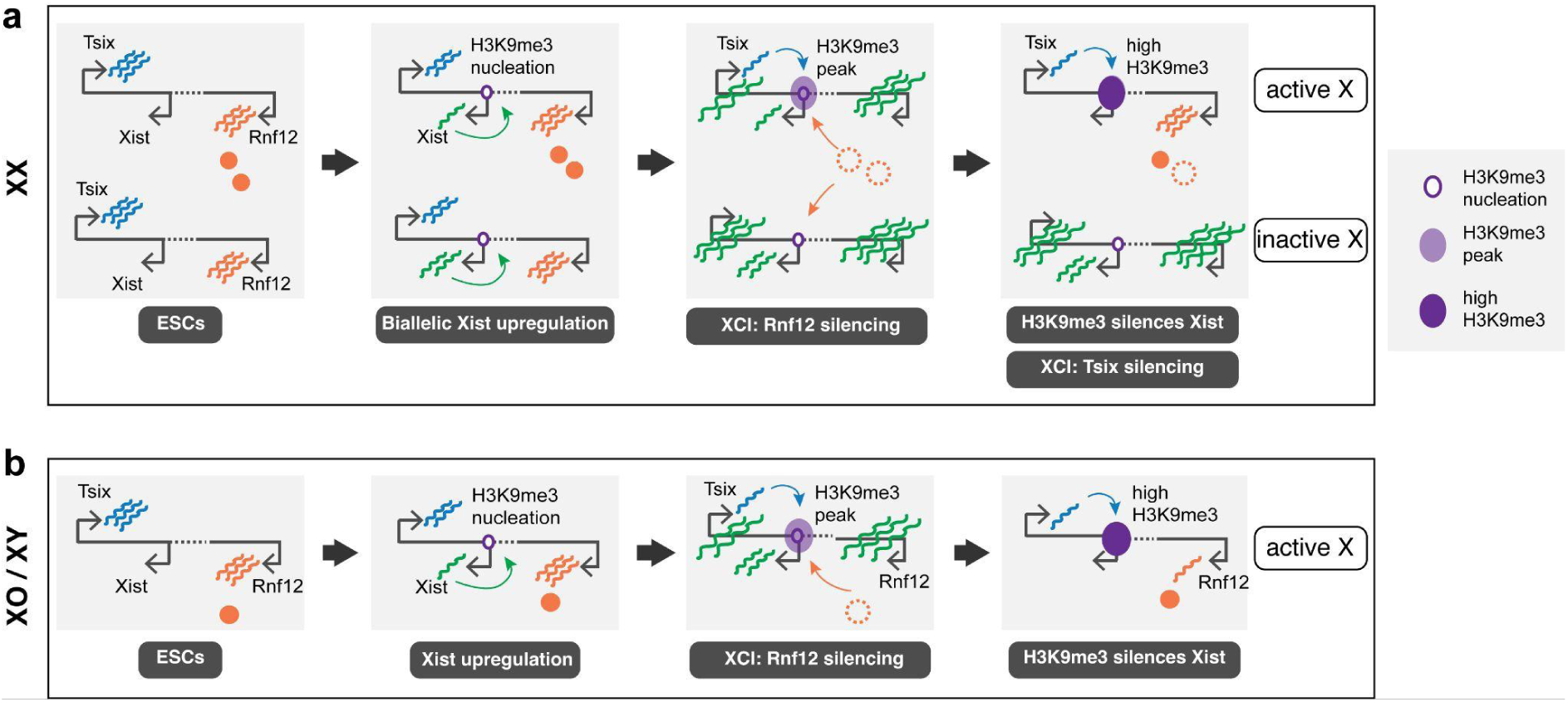
Xist-dependent heterochromatin shapes the random choice of the inactive X chromosome. Schematic illustration of the suggested model of XCI onset (a) In cells with two X chromosomes, biallelic Xist upregulation instructs H3K9me3 nucleation, which is further enhanced by Rnf12 silencing on both X chromosomes. Absence of RNF12 enforces Xist-dependent H3K9me3 accumulation around the *Xist* promoter, preferentially on the allele where Tsix is still transcribed. Rnf12 is silenced by XCI much faster than Tsix (Barros de Andrade E Sousa *et al*, 2019), therefore allowing a window in which both mechanisms can support H3K9me3 accumulation. Heterochromatin formation leads to downregulation of Xist and resolves the biallelic state into monoallelic expression. (b) In male cells, transient Xist upregulation nucleates H3K9me3 and leads to Rnf12 being silenced which, similarly to XX cells, enforces H3K9me3 accumulation around the *Xist* promoter. This leads to Xist downregulation and therefore correction of aberrant XCI.

A key finding here is that Xist transcription is required for full H3K9me3 deposition in XX and XO cells. Even in cells with a single X chromosome, where Xist transcription is generally low, a further reduction impairs proper heterochromatin formation, revealing for the first time a functional role for Xist transcription in cells with a single X. Importantly, only H3K9me3 was affected by the Xist perturbation, but not DNA methylation. This finding is consistent with a recent study reporting that Xist transcription can directly recruit H3K9me3 through a HUSH-mediated mechanism (Almeida *et al*, 2026).

Our two seemingly contradictory findings, that Xist transcription recruits H3K9me3 *in cis* and that H3K9me3 is found on the silent *Xist* allele, lead to the question of how the allelic distribution of H3K9me3 is controlled. A piece of the puzzle could be Tsix, which is silenced by highly transcribed Xist and thereby restricted to the *Xist*-silent allele. We show that Tsix is not required for Xist-mediated initiation of H3K9me3, but that it supports maintenance of the mark during differentiation. This could lead to preferential H3K9me3 maintenance on the silent *Xist* allele as follows: Assuming the presence of a heterochromatin nucleation signal (for example the stochastic, low Xist transcription), Tsix might promote spreading and accumulation of the mark through increasing DNA methylation levels via H3K36me3 deposition. Increased DNA methylation levels would in turn promote H3K9me3 maintenance through UHRF1, which binds hemimethylated DNA during replication and promotes recruitment of H3K9 methyltransferases (Tatarakis *et al*, 2025; Liu *et al*, 2025). Alternatively, Tsix might also directly promote H3K9me3 formation, for example through its previously reported recruitment of the HUSH-complex component MPP8 to the *Tsix* termination site (Spencley *et al*, 2023). Although Tsix is likely a contributing factor, the exact procedure of allele-specific H3K9me3 patterning has yet to be revealed. A key open question remains, why Xist transcription does not recruit higher H3K9me3 levels to the *Xist*-expressing allele, as observed in the Xist overexpression experiments.

We also tested known Xist regulators for a role in heterochromatin formation and found that RNF12 counteracts H3K9me3 and DNA methylation. RNF12 is a much characterized Xist activator and is generally thought to function by targeting the Xist repressor REX1 for degradation (Gontan *et al*, 2012, 2018). Here we have identified a second, REX1-independent pathway, which promotes Xist activation by counteracting H3K9me3 and DNA methylation. Our finding also explains why in a previous study a *Rex1* deletion could not fully rescue the Rnf12 KO phenotype (Gontan *et al*, 2018). Moreover, the REX1-independent pathway would enable RNF12 to contribute to proper Xist activation, even after the transcriptional downregulation of Rex1, which occurs shortly after cells exit the naive pluripotent state (Wang *et al*, 2023). The mechanism of the REX1-independent function remains an open question. The effect could be mediated by other global RNF12 targets, but also through ubiquitination of local chromatin components, since RNF12 has recently been found to directly bind to chromatin, including the RE57 element (Espejo-Serrano *et al*, 2024).

RNF12 is an Xist activator encoded on the X chromosome, and thereby part of a regulator class that has long been implicated in XCI regulation (Monkhorst *et al*, 2008). In a previous study using mathematical modelling we showed that such activators might control monoallelic Xist expression through two different principles: Their increased dose in female cells can promote female-specific Xist upregulation, and their absence upon silencing of too many X chromosomes can reverse erroneous Xist upregulation (Mutzel *et al*, 2019). In the latter case, silencing of the single X in males or of both X chromosomes in females will silence all copies of the activator, resulting in Xist downregulation. We propose that, in addition to contributing to female-specific Xist upregulation through the REX1 axis, RNF12 can also have a corrective role, that could be mediated by its effect on heterochromatin (Fig. 6): Absence of RNF12, when all X chromosomes in a cell are silenced, would result in heterochromatin formation at the *Xist* promoter region and “correct” spurious expression by silencing Xist. Such a mechanism might be even more important in species that exhibit higher rates of transient biallelic expression, such as primates and rabbits (Okamoto *et al*, 2011), as the need for correcting biallelic expression would be more pressing.

Random XCI in mice is typically framed as the decision of which *Xist* allele to upregulate. In other species, such as humans and rabbits, both *Xist* alleles are initially active, so choice is the decision of which *Xist* allele to downregulate. Our finding that H3K9me3 is involved in choice during mouse XCI points to a shared mechanistic basis across species: the choice of the *Xist*-silent allele might also occur in mice, but at faster timescales compared to human XCI. We therefore propose a model where transient, low Xist expression recruits H3K9me3 and marks the allele for silencing. Such a mechanistic coupling of activating and repressing mechanisms might provide a buffer time for the cell to ensure strictly monoallelic expression. Future work will hopefully map out how an initially active *Xist* allele is later silenced, and how this phenomenon relates to Xist levels. Lastly, future work is needed to show whether H3K9me3 has a role in establishment and maintenance of monoallelic Xist expression *in vivo*.

## Methods

### Cell lines

The female (XX) mouse embryonic stem cell line TX1072 (clone A3) is an F1 hybrid cross of C57BL/6 (B6) and CAST/EiJ (Cast) mouse strains (Schulz *et al*, 2014). The *Xist* promoter on the B6 allele is doxycycline-inducible, as there is a Tetracycline Response Element with seven copies of *tetO* followed by a minimal CMV promoter right upstream of the *Xist* transcription start site. The reverse tetracycline-controlled trans-activator (rtTA) is inserted in the Rosa26 locus.

The XO cell line (TX1072 XO clone B7) is derived from TX1072, with a loss of the B6 chromosome (Gjaltema *et al*, 2022). It is also trisomic for chr16. The XX_ΔXIC-B6_ (clone A1) and the XX_ΔXIC-Cast_ (clone A6) are derived from the TX1072 cell line with a deletion around the *Xist* locus on the B6 and the Cast allele, respectively (chrX:103,182,701 − 103,955,531 and chrX:103,183,145-103,955,698 mm10) (Pacini *et al*, 2021).

The female CasTuner cell line (TX SP427 clone B2) (Schwämmle *et al*, 2025) was derived from TX1072 and harbors a stable integration of the dCas9-HDAC4 fused with the conditional degron domain FKBP12^F36V^ (CasTuner system, (Addgene, 187956)) (Noviello *et al*, 2023). dTAG-13 (Tocris) addition to cell culture media (500 nM) ensures degradation of the CRISPRi system. The female split dCas9-KRAB cell line (TX SP107 cell line clone B6) has a stable integration of PYL1-KRAB-IRES-Blast and ABI-tagBFP-SpdCas9, that dimerize upon abscisic acid (ABA) addition to the cell culture medium (100 μM final concentration) (Gjaltema *et al*, 2022). All CRISPRi cell lines were transduced with plasmids carrying sgRNA sequences for the targeted genes (see Lentiviral transduction).

Female mouse ES cell line F121.9 (129/Sv-Cast/EiJ) and the derived deletion cell lines with a homozygous deletion of Rnf12 (RNF12-/- (Clone HG), in short Rnf12 KO) alone, or in addition to a Rex1 homozygous deletion (double knock-out RNF12-/-REX1-/- Clone #27.8, in short DKO), were a gift from Dr. Cristina Gontan Pardo (Gontan *et al*, 2018). In brief, Rnf12 KO was generated with CRISPR/Cas9-mediated removal of the full open reading frame of Rnf12, and the double knock-out was generated by additional CRISPR/Cas9-mediated deletion of most of the Rex1 open reading frame.

The XX status of all cell lines was tested before each experiment with RNA-FISH using probes for the X-linked gene *Huwe1* and for *Tsix* (Dxpas34 repeat), as previously described (Mutzel et al 2019).

### Cell culture and differentiation

TX1072 mouse ES cells and all derived cell lines were cultured in feeder-free 2i/FBS/LIF conditions (DMEM (Sigma-Aldrich D6429), 15% ES cell-grade foetal bovine serum (FBS, Pan Biotech P30-2602), 0.1 mM β-mercaptoethanol, 1000 U/mL LIF (Millipore), 3 mM GSK3 inhibitor CT-99021 (Axon Medchem) and 1 mM MEK inhibitor PD0325901 (Axon Medchem)). For CasTuner lines, 500nM dTAG-13 was added to the medium. They were plated at a density of 3.2 × 10^4^ cells per cm^2^ on gelatin-coated T25 flasks (Corning). The medium was changed daily, and the cells were split every two days.

HEK293T cells for lentivirus production were cultured on cell-culture plates in DMEM, 10% FBS (Gibco) and 0.1 mM β-mercaptoethanol.

WT F121.9 mouse ESCs and the derived deletion cell lines RNF12-/- and RNF12 -/- REX1 -/- (DKO) were adapted to feeder-free 2i/FBS/LIF conditions (DMEM (GIBCO 11995065), 15% ES cell-grade foetal bovine serum (FBS, Pan Biotech P30-2602), 100 U/mL penicillin/streptomycin (Thermo Fisher Scientifc), 1x NEAA (Gibco 11140035), 0.1 mM β-mercaptoethanol, 1000 U/mL LIF (Millipore), 3 mM GSK3 inhibitor CT-99021 (Axon Medchem) and 1 mM MEK inhibitor PD0325901 (Axon Medchem)) for 5 passages. They were plated at a density of 3.2 × 10^4^ cells per cm^2^ on gelatin-coated T25 plates (Corning). The medium was changed daily, and the cells were split every two days.

Differentiation of all cell lines was induced by 2i/LIF withdrawal (DMEM (Sigma-Aldrich), 10% FBS (Gibco) and 0.1 mM β-mercaptoethanol). The cells were plated at a density of 2.1 × 10^4^ cells per cm^2^, on 6-well plates coated with 10 μg/mL fibronectin (Corning 356008 or Serva 21370.03) in PBS for 1 hour. Once seeded for differentiation, the medium was changed every day and the cells were not split until harvesting.

### Generation of TX-Maged1-2tag ESCs

Two donor plasmids for Maged1 tagging were generated: SP756 containing the following construct: T2A-NLS-mStayGold-FKBP12^F36V,L106P^; and SP763 containing: T2A-NLS-mScarlet-I3-FKBP12^F36V,L106P^, flanked by the homology arms. FKBP12 is a destabilization domain carrying mutations F36V (sensitivity to dTAG-13), L106P (sensitivity to Shield1). As plasmid backbone the MTK0_017 plasmid, a gift from Hana El-Samad (Addgene plasmid # 123932 ; http://n2t.net/addgene:123932 ; RRID:Addgene_123932) (Fonseca *et al*, 2019), was used. After digestion with 10 units of Esp3I and 20 units of XhoI restriction enzymes (NEB), the plasmid backbone containing the Kanamycin resistance and the ColE1 (origin of replication) was purified from a 1% agarose gel. The plasmids were assembled by Golden Gate assembly, using the vector backbone and 4 additional fragments F1-F4. F1 contained the 5’ homology arm covering chrX:94,535,871-94,536,470 (600bp), F4 encompassed the 3’ homology arm spanning chrX:94,535,271-94,535,870 (600bp), F2 consisted of the T2A sequence followed by a nuclear localization sequence (NLS), the mScarlet-I3 or mStayGold sequence, a GGS linker and part of the FKBP12^F36V, L106P^ sequence (Supplementary Table 3) and F3 contained the rest of the FKBP12^F36V, L106P^ sequence (Supplementary Table 3). F1 and F4 were ordered as gene blocks (IDT), F2 and F3 were amplified by PCR from a plasmid. The guide plasmid was generated by cloning a sgRNA sequence targeting the C-terminus of the X-linked gene *Maged1* (sequence in Supplementary Table 2) into the pSpCas9(BB)-2A-Puro (PX459) V2.0, a gift from Feng Zhang (Addgene plasmid # 62988 ; http://n2t.net/addgene:62988 ; RRID:Addgene_62988) (Ran *et al*, 2013) containing the Cas9 sequence. PX459 was digested with 10 units of BbsI restriction enzyme (NEB). The digested DNA was purified from a 1% agarose gel and used as a vector to clone the guide by ligation using the T4 Ligase (NEB).

TX1072 cells were transfected by lipofection with three different plasmids: the guide plasmid and the two donor plasmids. To isolate cells that had integrated each construct in a different allele, cells showing both mScarlet-I3 and mStayGold signals were sorted by FACS two days after the transfection. A second sort to enrich the mStayGold^high^/mScarlet-I3^high^ cells was performed ten days after the first sort, followed by the generation of clonal cell lines. Two days before each sort, 1μM Shield1 (Takara) was added to the medium to stabilize the fluorescent proteins. The configuration of the allele-specific tagging was tested by Xist induction through the Dox-inducible promoter at the B6 allele. The clone E6 was chosen by genotyping PCRs and NGS karyotyping.

### Generation of TX XO CasTuner line

To knock down Xist in XO cells with CRISPRi, a monoclonal XO CasTuner cell line was created. In detail, TX1072 cells were transfected using Lipofectamine 3000 (Invitrogen) with the pSLPB2B-FKBP12_F36V-hHDAC4-SpdCas9-tagBFP-PGK-Blast plasmid (Addgene, 187956) (Noviello *et al*, 2023) and a vector carrying a hyperactive PiggyBac transposase (pBROAD3-hyPBase-IRES-zeocin) at a 5:1 molar ratio. Successful integrations were selected with blasticidin (5 μg/μL; Roth). The monoclonal A2 line was generated by fluorescence-activated cell sorting (FACS) based on high tagBFP expression and NGS karyotyping. As the parental line was XX, karyotyping and length-polymorphism PCR for the presence of X chromosomes (primers VM151 and VM152) revealed the A2 clone had lost the Cast allele but otherwise retained normal karyotype (39, XO).

### Lentiviral transduction

CRISPRi multiguide plasmids were stably integrated into mES cells via lentiviral transduction. For virus production, HEK293T cells were plated at 1.1 × 10⁵ cells/cm² (1 million cells in one well of a 6-well plate) and when reaching 95% confluency (the next day) they were transfected with a third-generation lentiviral system comprising VSVG, pLP1, and pLP2 (Thermo Fisher). Specifically, 1.2 μg pLP1, 0.6 μg pLP2, and 0.4 μg VSVG were combined with 2 μg multiguide plasmid in 250 μL OptiMEM and incubated for 5 min. In parallel, 11.25 μL Lipofectamine 2000 was incubated in 250 μL OptiMEM for 5 min. Then, the two mixtures were combined and incubated for another 15 min, before being added to the HEK293T cells. The supernatant containing the virus was collected 48 h later and concentrated tenfold using the Lenti-X concentrator (Takara).

The ESCs to be transduced were cultured in serum/LIF conditions to prevent karyotype abnormalities (loss of one X chromosome), and were plated one day prior to transduction (0.2 million cells in a 12-well). The day of the transduction, 8 ng/μL polybrene (Merck) was added to the culture media to improve transduction efficiency, and between 50 and 100 μL of concentrated virus was used per transduction. Forty-eight hours post-transduction, cells were selected for successful plasmid integration with puromycin (1 μg/ml; Sigma). Cells were subsequently cultured in 2i/FBS/LIF medium for at least five passages before any experiments.

### NGS Karyotyping

For karyotyping, shallow whole-genome sequencing was performed. For TX-*Maged1*-2tag ESCs we used the kit ExpressPlex™ 2.0 Library Prep Kit – 96-well for Illumina ® Sequencing Platforms by seqWell. For TX XO CasTuner ESCs, the karyotyping was performed via double digest genotyping-by-sequencing (ddGBS), as described in (Gjaltema *et al*, 2022). Briefly, the forward and reverse strands of a barcode adaptor and common adaptor were diluted and annealed, and they were pipetted into each well of a 96-well PCR plate together with 1 mg of each sample and dried overnight (oligo sequences are listed in Suppl. Table 2). The following day, the samples were digested with 20 mL of a NIaIII and PstI enzyme mix (New England Biolabs) in NEB Cutsmart Buffer at 37°C for 2 h. After the digest, a 30 mL mix with 1.6 mL of T4 DNA ligase (New England Biolabs) was added to each well and placed on a thermocycler (16°C 60 min followed by 80°C 30 min for enzyme inactivation) to ligate to the genomic DNA the adapters with ends complementary to those generated by the two restriction enzymes. Samples were cleaned with CleanNGS beads (CleanNA) using 90 mL of beads for each well and following manufacturer’s instructions. Samples were eluted in 25 mL ddH2O and DNA was quantified using a dsDNA HS Qubit assay (Thermofisher). Samples were pooled in an equimolar fashion, size-selected (300-450bp) by loading 400 ng of each pooled sample on an agarose gel and cleaned the Nucleospin Gel and PCR Cleanup kit (Macherey-Nagel). Samples were PCR amplified using the Phusion High-Fidelity DNA Polymerase (New England Biolabs) and an annealing temperature of 68°C over 15 amplification cycles (OG218/OG219, Suppl. Table 2). Resulting amplicons were cleaned with CleanNGS beads in a 1:1.2 ratio (sample:beads) and sequenced.

### NGS Karyotyping analysis

Sequencing reads were mapped to the mouse genome (mm10) with bowtie2 (Langmead & Salzberg, 2012). Unmapped reads and reads of low (<20) quality were discarded with samtools view [-F 4 -q 20] and the file was sorted and indexed with samtools. Reads per chromosome were counted with deepTools multiBamSummary (Ramírez *et al*, 2016).

### FACS

Fluorescence-activated cell sorting (FACS) was used in the creation of the TX-*Maged1*-2tag cell line, to select for cells that have successfully integrated the two tags, and later on for the experiment of sorting monoallelic Xist-expressing populations. A BD FACS Aria II flow cytometer (BD Biosciences) with a 100μm nozzle and a 5B-3YG-2R-1DB laser configuration was used. BD FACS Diva software (v8.0.1) was used for setting the gates. The cells were harvested and resuspended in FACS buffer (PBS, 10% ESC-grade FBS (Gibco), 0.5 mM EDTA) on ice. They were then sorted based on high mScarlet-I3 (615/20) and mStayGold (510/20) levels. An example of gating strategy is shown in (Suppl. Fig. 2g). The sideward and forward scatter areas were used for live cell gating. The height and width of the forward scatter and subsequently the height and width of the sideward scatter were used for singlet/doublet differentiation.

Sorted cells were centrifuged, resuspended in cell culture medium, and if re-seeded the medium was supplemented with 1x Penicillin-Streptomycin (Thermo Fisher Scientific) for 10 days.

Gates from BD FACS Diva software were loaded in R (v4.5.2) with packages flowWorkspace (v4.22.1) and CytoML (v2.22.0), and visualized with ggcyto (v1.38.1).

### CRISPR inhibition

For the knock-down of Zfp36l1 and Zfp280c, the CasTuner system was induced with dTAG-13 removal for 6 days in total, the last 4 of which differentiation was also induced. The control condition was cultured with dTAG-13 and differentiated for 4 days. To knock-down the transcription of Rex1 and Zfp281, the dCas9-KRAB system was induced with ABA for 5 days in total, the last 4 of which differentiation was also induced. The safe-targeting control line with “STC” guides was cultured in the same conditions as the knock-down cell lines.

To knock-down the transcription of Rnf12 and Zic3, the CasTuner system was induced by dTAG-13 removal for 7 days in total, and during the last 3 days differentiation was induced. The control condition was cultured with dTAG-13 and differentiated for 3 days.

The knock-down of Xist (in TX1072 XX and XO) and Tsix (in TX1072 XX) with CasTuner was induced with dTAG-13 removal from the culture medium for 6 days in total, and differentiation was induced during the last 4 days before harvesting. The knock-down of Tsix in the experiment of parallel Xist overexpression was induced for 4 days in total, without simultaneous differentiation.

Xist overexpression was induced with Doxycycline addition (1µg/ml) for two days, with and without differentiation. For transient Xist overexpression, Doxycycline was added for one or two days, and then washed out.

### Multiguide plasmids

To target the CRISPRi system to the promoter of the desired gene, sgRNA sequences were designed with Guidescan2 or Chopchop v3 (Labun *et al*, 2019; Schmidt *et al*, 2025). Three to four different sgRNA sequences were cloned into the SP199 sgRNA expression plasmid using Golden Gate assembly. A distinct Pol III promoter (hu6, mu6, hH1, or 7sk) controls the expression of each guide. Promoter–sgRNA-handle constructs were PCR-amplified with primers containing the guide sequences and a BsmBI/Esp3I site, then ligated into Esp3I-digested SP199 using T4 ligase and Esp3I (New England Biolabs). Golden Gate reactions were performed for 20 cycles (5 min at 37 °C, 5 min at 20 °C) followed by a 65 °C incubation for 20 min. Constructs were transformed into NEB Stable *E. coli* and verified by ApaI digestion and Sanger sequencing. Guide sequences are provided in Suppl. Table. 2.

### RNA-FISH

Cells were harvested with accutase (Thermo Fisher Scientific) for 5 min, resuspended in medium, then adhered on Poly-L-Lysine (Sigma Aldrich) coated glass coverslips (10 min incubation) and fixed with 3% paraformaldehyde for 10 min. After washing with PBS, cells were permeabilized for 5 min with 1xPBS, 2 mM Ribonucleoside Vanadyl Complex (NEB), and 0.5% TritonX-100 (Sigma Aldrich), and then stored in 70% ethanol at -20°C.

SNP-FISH was developed to distinguish between Xist RNA molecules expressed from the B6 (Q570/Atto550) or the Cast (Q670/Cy5) allele. Transcripts were detected with 28 Stellaris probes (Biosearch Technologies), each targeting a SNP (Supplementary table 2).

To allow specific binding to one sequence variant, part of the probe is initially masked through a complementary shorter oligo. To this end, for each allele, the Stellaris SNP oligos (12.5µM) were annealed to shorter mask oligos (12.5µM), by mixing them 1:1 and heating the mix at 75°C for 2 min, ramping down to 25°C by 0.1°C per sec (ramp rate 5%), holding at 25°C for 30 min, and keeping at 4°C. The annealed oligos were used immediately for hybridization.

In conventional RNA-FISH, Xist transcripts were detected with exonic Stellaris probes (Biosearch Technologies). In both RNA-FISH and SNP-FISH, fixed cells were hybridized overnight at 37 °C with 250 nM of each FISH probe (exonic probes for RNA-FISH, B6 and Cast probes for SNP-FISH) in 50 μL of Stellaris RNA-FISH hybridization buffer (Biosearch Technologies) containing 10% formamide. Coverslips were washed with Wash buffer A (2× SSC buffer (Sigma-Aldrich) and 10% formamide in water), for 30 min at 37 °C. Then they were transferred to Wash buffer A containing 0.2 mg/mL DAPI (Sigma-Aldrich) for 30 min at 37 °C. They were washed with 2xSSC in water at room temperature for 5 min, and mounted on glass slides with Vectashield (Biozol). FISH probe sequences are provided in Suppl. Table 2.

Images were acquired with a Z1 Observer microscope (Zeiss) using a ×100 objective. Xist clouds were counted manually for at least 100 cells per sample.

### Real-time qPCR

Cells were lysed with Trizol (Invitrogen), and RNA was extracted with Direct-zol RNA purification kit (Zymo Research). Reverse-transcription of RNA into complementary DNA (cDNA) was performed with Superscript III reverse transcriptase (Invitrogen), using random hexamer primers. The cDNA was then used as template for real-time qPCR with Power SYBR Green PCR master mix (Thermo Fisher Scientific) and gene-specific primers (Suppl. Table 2). The qPCR reaction was carried out in Quant-Studio 7 Flex real-time PCR machine (Thermo Fisher Scientific) and quantified in R. Relative gene expression was calculated compared to the geometric mean of the housekeeping genes Rrm2 and Rplp0.

### CUT&Tag

CUT&Tag (Kaya-Okur *et al*, 2019) was performed on cells directly after harvesting. Cells were dissociated with accutase for 5 min, followed by media addition to stop dissociation. 10^5^ cells were collected per CUT&Tag reaction and washed with 1mL wash buffer (20 mM HEPES–KOH pH 7.5, 150 mM NaCl, 0.5 mM spermidine, 10 mM sodium butyrate, protease Inhibitor and 1 mM PMSF). After washing, cells were resuspended in 100 μL wash buffer. For every 10^5^ cells, 10 μL of BioMag Plus Concanavalin A beads were used, after being equilibrated in excess of binding buffer (20 mM HEPES–KOH pH 7.5,10 mM KCl,1 mM CaCl_2_, 1 mM MnCl_2_) and concentrated in 10 μL of binding buffer. The cells were incubated with Concanavalin A beads by rotating for 10 min at room temperature. The beads with the bound cells were separated with a magnet and resuspended in 100 μL of ice-cold antibody buffer (wash buffer with 0.05% digitonin and 2mM EDTA). 1 μL of primary antibody (H3K9me3, Active Motif #39161) was added and beads were incubated on a rotator for 3 h at 4 °C. The beads were magnetically separated and resuspended in 100 μL of ice-cold Dig-wash buffer (wash buffer with 0.05% digitonin) and 1 μL of secondary antibody (Guinea pig anti-rabbit, antibodies online ABIN101961) and incubated for 1 h at 4 °C on a rotator. Next, the beads were washed three times with ice-cold Dig-wash buffer, and resuspended in 100 μL ice-cold Dig-300 buffer (20 mM HEPES–KOH pH 7.5, 300 mM NaCl, 0.5 mM spermidine, 0.01% digitonin, 10 mM sodium butyrate and 1 mM PMSF and protease inhibitor) with 0.4 μL 3×FLAG–pA-Tn5 (5.5μM) (derived from Addgene 124601 and purified according to (Gjaltema *et al*, 2022)) preloaded with mosaic-end adaptors. Beads were incubated for 1 h at 4 °C on a rotator, and then washed four times with ice-cold Dig-300 buffer. While on ice, beads were resuspended in 50 μL of tagmentation buffer (Dig-300 buffer, 10 mM MgCl_2_ and 0.125pM spike-in DNA). Spike-in DNA was prepared with PCR amplification of the Ampicillin resistance gene (AmpR) that also contains i5 and i7 primer binding sites on 5′ and 3′ ends, respectively (Supplementary Table 2). Tagmentation was carried out for 1 h at 37 °C. To stop tagmentation and solubilize DNA fragments, 2.25 μL of 0.5 M EDTA, 2.75 μL of 10% SDS and 0.5 μL of 20 mg/mL proteinase K were added, followed by vortexing for 5 s. Samples were incubated overnight at 55 °C followed by 30 min at 70 °C to inactivate proteinase K. The DNA fragments were purified with the ChIP DNA clean and concentrator kit (Zymo Research) and eluted in 25 μL of elution buffer. Libraries for sequencing were prepared by amplifying the purified fragments with indexed primers and NEBNext HiFi 2× PCR Master Mix for 14 amplification cycles. Post-PCR cleanup was performed with AMPure XP beads (1x volume), and the final libraries were eluted in 27 μL of 10 mM Tris (pH 8.0).

Libraries were sequenced 2x75bp or 2x150bp paired-end with NextSeq 2000, NovaSeq 6000, DNBSEQ G400 or Aviti sequencers.

### CUT&Tag analysis

Sequencing reads were trimmed with Trim Galore 0.6.10. Quality controls were performed before and after trimming with FastQC v0.12.1 and multiqc, version 1.26 (Ewels *et al*, 2016). Reads were mapped to mm10 using bowtie2 version 2.5.0 (Langmead & Salzberg, 2012). with settings [--end-to-end --very-sensitive --no-mixed --no-discordant --phred33 -I 10 -X 2000].

Unmapped reads were filtered out using samtools (v1.22) view [-F 0x04]. ENCODE Blacklisted regions (mm10-v2) (Amemiya *et al*, 2019) were removed using bedtools intersect [-v] (v2.30.0). Coverage bigwig tracks were generated from filtered and sorted bam files using bamCoverage of the deepTools package (v3.5.6) (Ramírez *et al*, 2016), using [--extendReads -bs 1 --normalizeUsing RPKM].

Spike-in reads were not used for normalization, but were instead compared across conditions to ensure that the perturbations did not cause global changes in H3K9me3 levels.

Quantification of CUT&Tag signal over RE57 (chrX:103,481,046-103,482,684, mm10) was computed with bigWigAverageOverBed.

For visualization, bigwig files of replicates were merged with wiggletools mean and resulting wig files converted to bigwig with wigToBigWig.

To assign allelic reads, samtools mpileup was used (Li *et al*, 2009), supplying a bed file with SNP positions in a given range including RE57 and ∼2kb downstream (chrX:103,479,041-103,482,895). The contribution of each allele as a percentage of total reads was calculated in R, by counting reference and alternative alleles.

Peaks were called in each condition and replicate separately using epic2 (Stovner & Sætrom, 2019) with settings [--keep-duplicates --gaps-allowed 1]. Differential peak analysis was performed with DiffBind 3.16.0 in R version 4.4.2. Specifically, the consensus peak list was created from peaks found in at least 3 samples with dba.count settings as [minOverlap = 3, filter=5, bUseSummarizeOverlaps=TRUE, summits=FALSE, score=DBA_SCORE_RPKM] and normalization with [normalize = DBA_NORM_LIB, library = DBA_LIBSIZE_FULL]. Peaks with DESEQ2 FDR <0.05 are considered differential.

### Amplicon bisulfite sequencing

Genomic DNA was extracted from harvested cells using the DNeasy Blood & Tissue kit (Qiagen). 1μg of DNA were bisulfite-converted with EZ DNA Methylation-Gold kit (Zymo). One fifth of the eluted DNA was used per amplicon PCR. Primers for bisulfite-converted DNA targeting the *Xist*-promoter proximal CpG island (chrX:103,482,090-103,482,350 mm10 from UCSC Genome Browser) were designed with MethPrimer2.0 (Li & Dahiya, 2002). Sequences are provided in Suppl. Table 2. In total, five amplicons were designed, of which only the ones that target RE57 are included in this study. The other three amplicons cover the promoter of Xist (chrX:103,483,289-103,483,491 and chrX:103,483,101-103,483,344), and Major Satellite Repeats. PCR was performed with EpiTaq polymerase (Takara), using 25mM MgCl_2_. DNA was amplified for 27 cycles, at an annealing temperature of 53°C. PCR reactions of each sample were pooled (total volume for *Xist* amplicons) and cleaned with MinElute Reaction Cleanup kit (Qiagen) (2 columns per sample). Libraries were prepared with NEBNext Ultra II DNA Library Prep kit (New England Biolabs), indexed with NEBNext Multiplex Oligos for Illumina Dual Index Set 1 (New England Biolabs) and cleaned with AMPure XP beads (Beckman Coulter). Concentration and size distribution of libraries were analyzed with Qubit 1x dsDNA HS (Thermo Fisher Scientific) and HS DNA D1000 Tapestation (Agilent), respectively. Libraries were sequenced 2x150bp paired-end with MiSeq, DNBSEQ G400 or Aviti sequencers.

### Amplicon bisulfite sequencing analysis

Sequencing reads were trimmed with Trim Galore, with [--clip_R1 10 --three_prime_clip_R1 5 --clip_R2 15 --three_prime_clip_R2 5 --paired]. They were mapped using bsmapz with settings -m 0 -g 3 -n 1 -v 0.1 -w 10 to the reference fasta file (Supplementary Table 3) that contains the amplicon sequences of all amplicons used in the experiment, that is RE57, the *Xist* promoter and annotated Major Satellite Repeats. bam files were sorted and indexed with samtools, and were used for methylation calling by mcall command of MOABS package (Sun *et al*, 2014) [--trimWGBSEndRepairPE2Seq 0 --trimWGBSEndRepairPE1Seq 0--reportCHX 0]. For plotting, a coverage cutoff of totalC > 10 for *Xist* amplicons was imposed. Name-sorted bam files were used with epialleleR 1.14.0 (Nikolaienko *et al*, 2022) to extract DNA methylation epialleles.

### Pyrosequencing

To quantify relative allelic expression for Xist, Rnf12 and Fam122b, an amplicon containing a SNP was amplified by PCR from cDNA using GoTaq G2 Hot Start Taq (Promega) with 2.5 mM MgCl2 for 40 cycles, at an annealing temperature of 58°C. The PCR product was sequenced using the Pyromark Q24 system (Qiagen). Primers are provided in Supplementary Table 2.

### WGBS

Whole Genome Bisulfite Sequencing (WGBS) for XO and XX_ΔXIC-B6_ was performed in parallel, as described in (Mutzel *et al*, 2025). XX_ΔXIC-B6_ data were published in (Mutzel *et al*, 2025). Briefly, TX XO B7 and XX_ΔXIC-B6_ mES cells were differentiated for 0, 2 or 4 days and genomic DNA was extracted with the DNeasy Blood and Tissue Kit (Qiagen). Bisulfite-converted DNA libraries were prepared using the Accel-NGS Methyl-Seq DNA library kit (SWIFT BIOSCIENCES). Specifically, 200ng (in 50 μL lowTE) of purified DNA were fragmented to ∼350 bp using the Covaris S2 system (10% duty cycle, intensity 5 for 2 × 45 s in Covaris AFA tubes) followed by a concentration step with Zymo columns (DNA Clean & Concentrator). 20 μL of DNA were used for bisulfite conversion overnight with the Zymo EZ-DNA methylation Gold Kit. Bisulfite-converted DNA was fully denatured by 2 min incubation at 95 °C and immediately transferred to ice. To anneal a truncated adapter, 15 μL of denatured DNA was mixed with 25 μL of Adaptase reaction mix (SWIFT) and incubated for 15 min at 37 °C, 2 min at 95 °C, and then cooled to 4 °C. Extension and second strand synthesis by using a primer complementary to the truncated adapter was performed mixing the Adaptase reaction mix with 44 μL of extension reaction mix (SWIFT) and incubation at 98 °C for 1 min, 62 °C for 2 min, 65 °C for 5 min and cooling to 4 °C. The samples were cleaned with 1.2 volumes of Ampure XP beads, 80% ethanol and eluted in 15 μL. Ligation of the second (truncated) adapter was performed at 25 °C for 15 min followed by an additional bead clean up with 1 volume of AmpureXP beads and 80% ethanol. Samples were individually indexed using unique dual indexed primer sets with 6 cycles of the following PCR program: 30s @ 98 °C, 6 cycles of 15s @ 98 °C, 30s @ 60 °C, 60s @ 68 °C, followed by a final 5 min incubation step at 68 °C. Libraries were cleaned up with 1 volume of Ampure XP beads. Quality was assessed using Agilent’s Bioanalyzer and concentration was determined by qPCR. Libraries were pooled equimolarly and sequenced on a NovaSeq 6000 S2 flowcell (Illumina) in paired end 150 mode to yield ∼ 120-278mio fragments.

### WGBS analysis

WGBS analysis was performed as described in (Mutzel *et al*, 2025). Briefly, adaptor and quality trimming of fastq files was performed with Trim Galore (v0.6.4) with the options [--clip_R1 10 --three_prime_clip_R1 5 --clip_R2 15 --three_prime_clip_R2 5 --paired]. Reads were aligned to the mouse genome (mm10) with BSMAPz [-q 20 -u -w 100]. The resulting bam files were sorted and low quality mapped reads were removed using samtools [view -q 10] (v1.10) (Li *et al*, 2009). Next, duplicate reads were removed with Picard (v2.7.1) using the options [MarkDuplicates REMOVE_DUPLICATES = TRUE]. Replicate bam files were merged and indexed using samtools. To obtain bedGraph files containing per-base CpG methylation metrics, MethylDackel (v0.3.0) was applied [extract --mergeContext --minDepth 2]. Mitochondrial regions were subsequently removed as well as blacklisted regions for mm10 using bedtools [intersect -v] (v2.29.2) (Quinlan & Hall, 2010). To obtain bigwig files for genome browser visualization, bedGraph files were converted using the UCSC software bedGraphToBigWig (v4). The mean DNA methylation of CpGs in RE57 (Suppl. Fig. 1c) was calculated with getMeth from the R package bsseq (Hansen *et al*, 2012).

## Data availability

H3K9me3 XY ChIP-seq tracks (Bleckwehl *et al*, 2021)H3K9me3 CUT&Tag in differentiating XX_ΔXIC-B6_ and XO cells (Gjaltema *et al*, 2022)in differentiating XX_ΔXIC-B6(Gjaltema *et al, 2022)*_from (Mutzel *et al*, 2025; Schwämmle *et al*, 2025) (Mutzel *et al*, 2025)H3K9me3 ChIP-seq from mouse embryo (Wang *et al*, 2018) was accessed from GEO: GSE97778. Sequencing data generated in this study was deposited at GEO: GSE339369.

## Acknowledgements

We would like to thank Dr. Cristina Gontan Pardo for sharing the WT F121.9, Rnf12 KO and Rnf12/Rex1 DKO cell lines; Melanie Piedavent-Salomon and Adeline Dehlinger for setting up the gating strategy and sorting the cells. We also thank the sequencing facility and IT service at the Max Planck Institute for Molecular Genetics for support, and the members of the Schulz lab for critically reading the manuscript. This work was supported by the Max-Planck Lise Meitner Excellence program, ERC Starting Grant CisTune (948771) and by the DFG priority program SPP 2202 (508000619) to E.G.S. T.S. was supported by the DFG (GRK1772, IRTG 2403). G.N. was supported by the European Union’s Horizon 2020 Research and Innovation Program (Marie Skłodowska-Curie ITN PEP-NET).

## Author contributions

E.K. and E.G.S. conceived the project and designed the experiments. I.P.C created the TX-Maged1-2tag cell line and performed the associated experiments. G.N. generated the TX XO CasTuner cell line, and performed cell culture for the dense DNA methylation time course. G.N., T.S. and E.K. generated gene-targeting CRISPRi cell lines. I.D. performed karyotyping of all cell lines, pyrosequencing and SNP-FISH. E.K. and I.D. and G.N. performed KD experiments. E.K. and L.M. performed aBS. R.A.F.G. performed WGBS, and M.B. analyzed it. E.K. performed all other experiments. E.K. and E.G.S. wrote the manuscript with input from all authors.

## Supplementary Material

**Supplementary Table 1:** differential peak analysis on H3K9me3 CUT&Tag from the following experiments: Rnf12, Zic3, Zfp280c, Zfp281, Rex1, Zfp36l1 KDs and Rnf12 KO and DKO.

**Supplementary Table 2:** Experimental materials (sgRNAs, primers, probes).

**Supplementary Table 3:** DNA sequences (TX-Maged1-2tag cloning, reference genome file for amplicon bisulfite sequencing analysis).

## Supplementary Figures

**Supplementary Figure 1:**
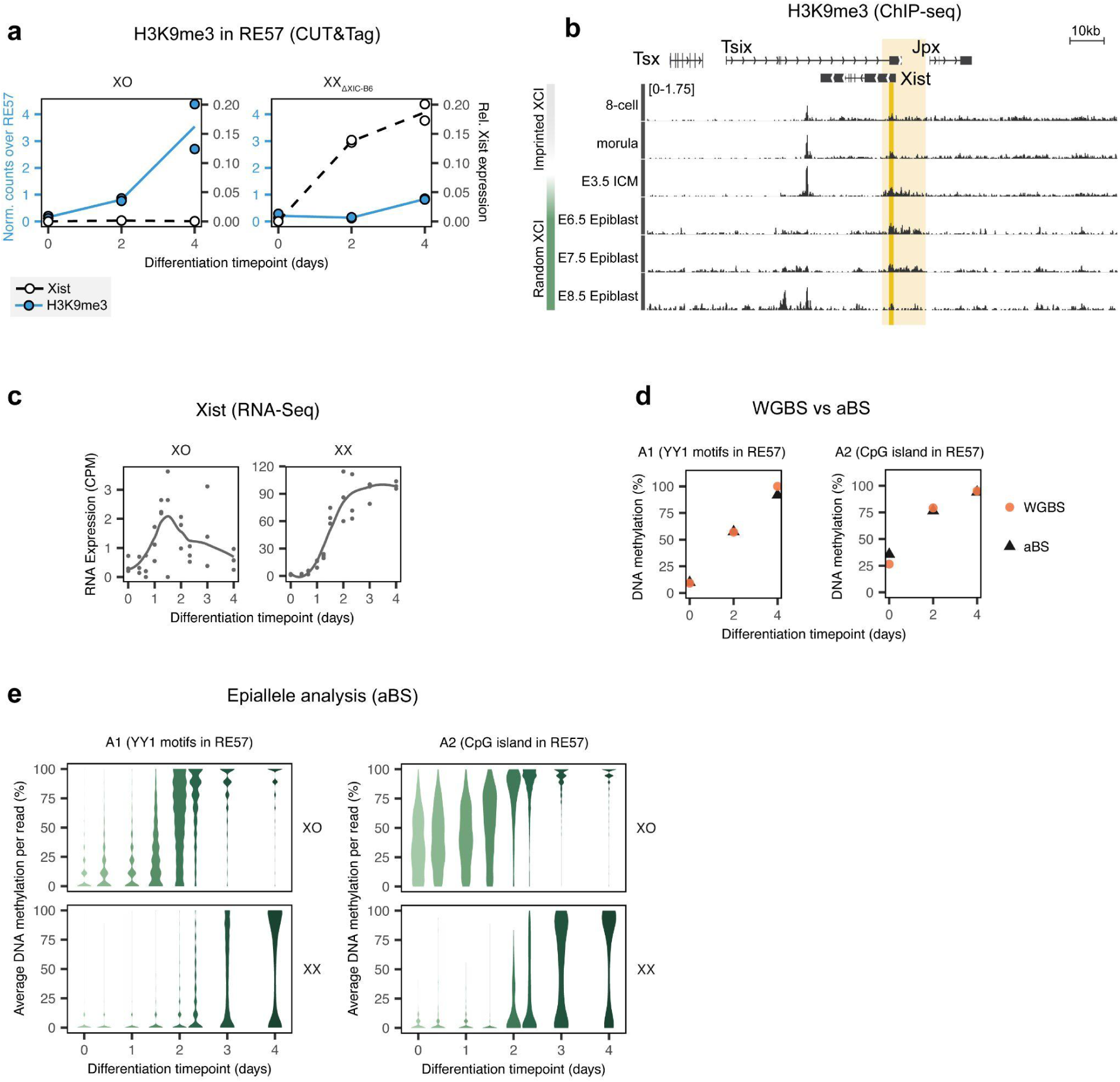
(a) Quantification of H3K9me3 from Fig. 1a in RE57 for XO and XX_ΔXIC-B6_ (blue) and Xist expression (RT-qPCR) in the same experiment (white marks). Data from (Gjaltema *et al*, 2022). (b) H3K9me3 ChIP-seq from mouse embryo data, average of two biological replicates (Wang *et al*, 2018). Light yellow box: H3K9me3-marked area around the *Xist* promoter, same region as Fig. 1a. Yellow box: RE57. Region shown: chrX:100,602,684 - 100,745,898 (mm9). (c) Xist expression dynamics in differentiation time course, RNA-seq (CPM) of three biological replicates (Schwämmle *et al*, 2025). The line represents a smoothed average of three replicates across all timepoints. (d) Comparison of DNA methylation in the XO cell line at covered amplicons in WGBS (mean of 2 biological replicates) and aBS (mean of 3 biological replicates). The two amplicons, A1 and A2, are shown, covering the YY1 binding sites and the CpG island in RE57 respectively. (e) Epiallele analysis of aBS data shows the distribution of average DNA methylation of individual DNA molecules. The violins have a fixed maximal width.

**Supplementary Figure 2:**
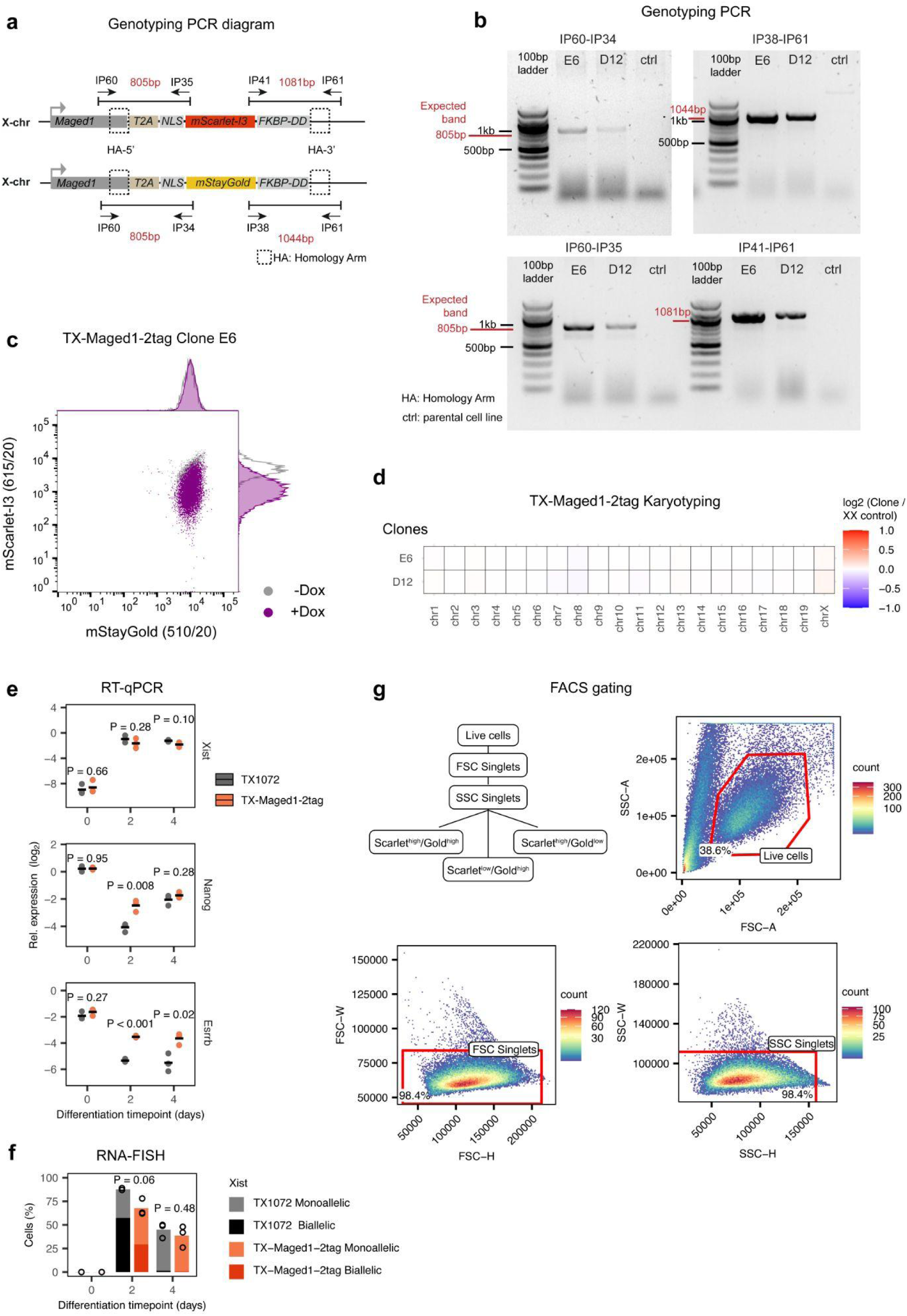
(a) Design of genotyping PCRs for TX-Maged1-2tag cell line. Primer positions are indicated as arrows. (b) Agarose gel images of genotyping PCRs for TX-Maged1-2tag clones D12, E6 and the parental cell line (TX1072 XX clone A3) as control. (c) Doxycycline treatment of TX-Maged1-2tag cell line clone E6 to determine on which allele each reporter is integrated: As the Dox-inducible *Xist* promoter is integrated on the B6 allele, Dox induction will inactivate the B6 X chromosome. Dox-treated cells (purple) show reduction in mScarlet-I3 signal and no change on the mStayGold signal compared to the non-treated cells (gray) by flow cytometry, indicating that mScarlet-I3 is integrated in the B6 allele. (d) Heatmap of Next Generation Sequencing (NGS) karyotyping data for the two clones of TX-Maged1-2tag. Counts mapping to each chromosome were normalized to the parental control cell line TX1072. (e) RT-qPCR for Xist and pluripotency markers for TX-Maged1-2tag and the parental cell line TX1072. P-value shown for two-sided, unpaired T-test. (f) RNA-FISH for differentiation time course of TX-Maged1-2tag and parental TX1072 cell line. P-value of two-sided unpaired T-test of Xist-positive cells between the two cell lines is shown. Dark colors: biallelic cells, light colors: monoallelic cells. (g) FACS gating strategy for sorting cell populations at differentiation day 4, used in main Fig. 2b. One representative biological replicate shown. Percentage of cells inside each gate are indicated. SSC: side scatter, FSC: forward scatter, A: area, H: height, W: width.

**Supplementary Figure 3:**
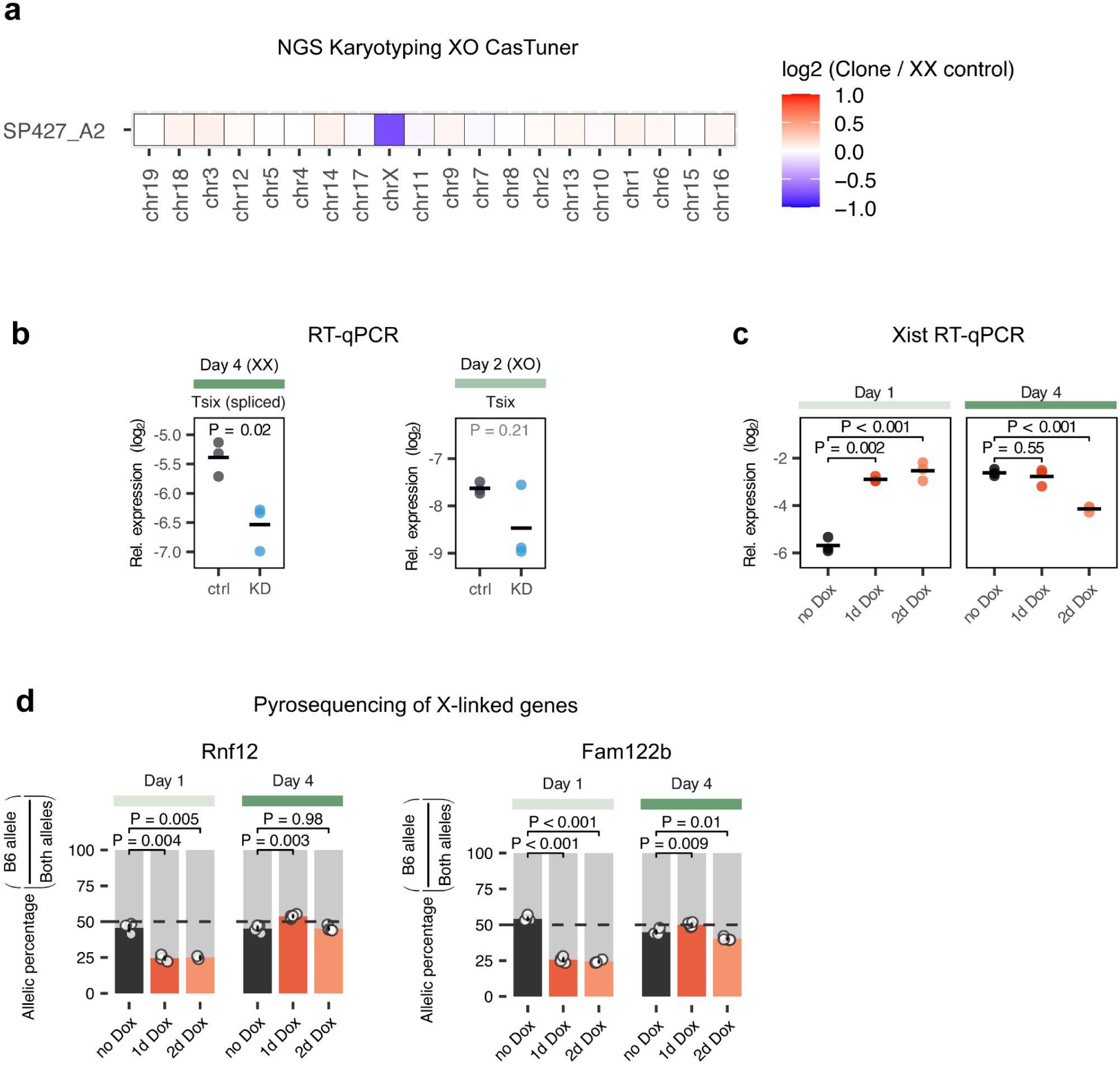
(a) NGS karyotyping for XO CasTuner cell line Clone A2, using TX1072 XX ESCs as control. (b) RT-qPCR data for Tsix expression. Horizontal lines indicate the mean of 3 biological replicates (dots). ctrl: +dTAG, KD: -dTAG. Two-sided unpaired T-test, p-value black if <0.5. (c) RT-qPCR for Xist at transient Xist overexpression experiment. Horizontal lines represent mean of three biological replicates (dots). P-value is reported for two-sided unpaired T-test. (d) Pyrosequencing of Rnf12 and Fam122b RNA. Percentage of reads coming from the B6 allele at each differentiation day and treatment are plotted. Error bar shows standard deviation. Two-sided unpaired T-test. Between 3 and 4 replicates for each condition.

**Supplementary Figure 4:**
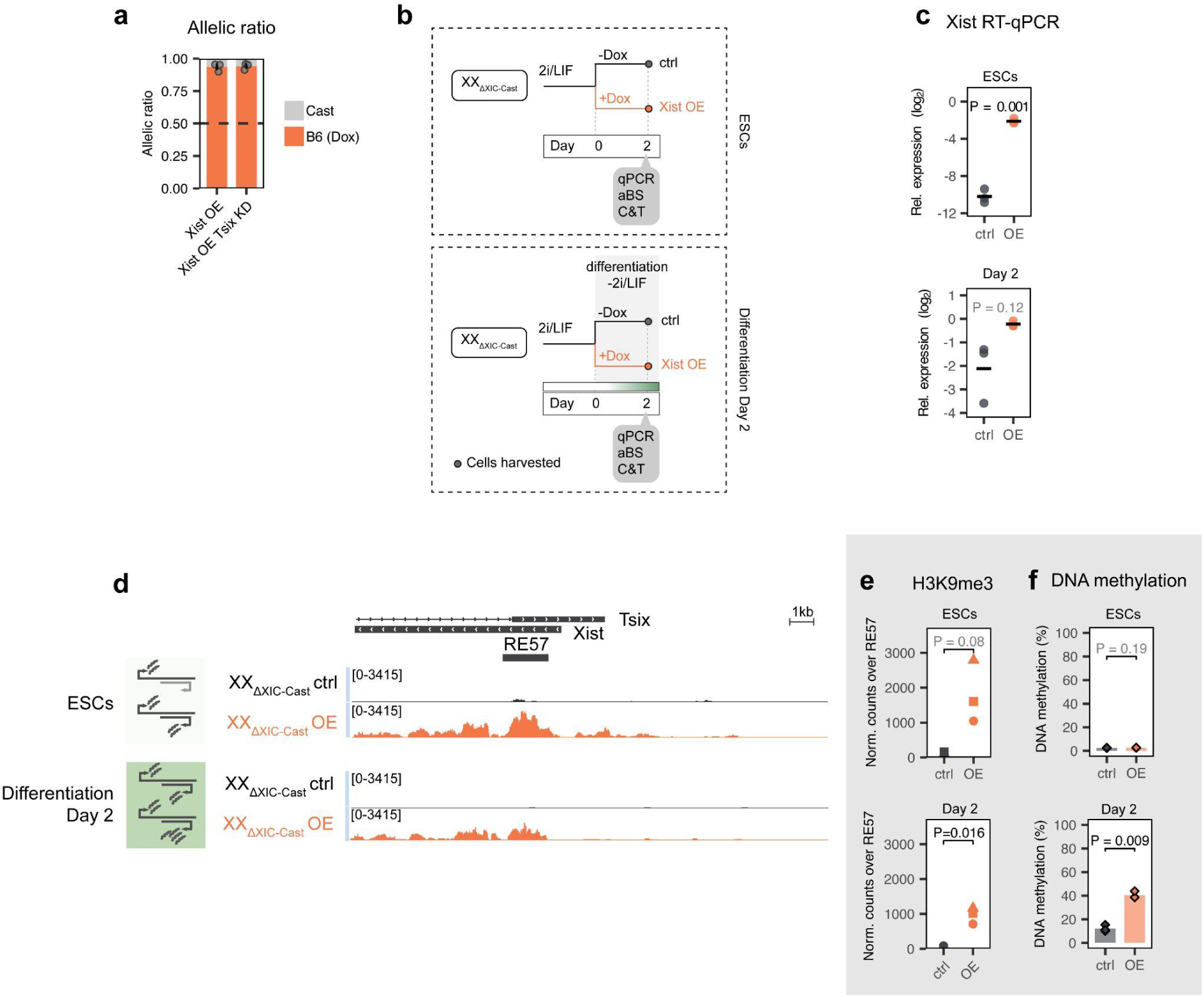
(a) Ratio of H3K9me3 CUT&Tag reads from the Dox-induced allele spanning extended RE57 region (chrX:103,479,041-103,482,895). (b) Experiment outline for Xist overexpression in XX_ΔXIC-Cast_ in undifferentiated ESCs and during differentiation (day 2), used in c-f. (c) RT-qPCR for Xist. Horizontal lines represent the mean of three biological replicates (dots). Two-sided unpaired T-test. (d) H3K9me3 CUT&Tag tracks (average of 3 biological replicates, normalized bigwig). (e) Normalized H3K9me3 reads in RE57. Two-sided paired T-test. (f) DNA methylation measured by aBS. The bars represent the mean of 3 biological replicates (diamonds). Two-sided paired T-test.

**Supplementary Figure 5:**
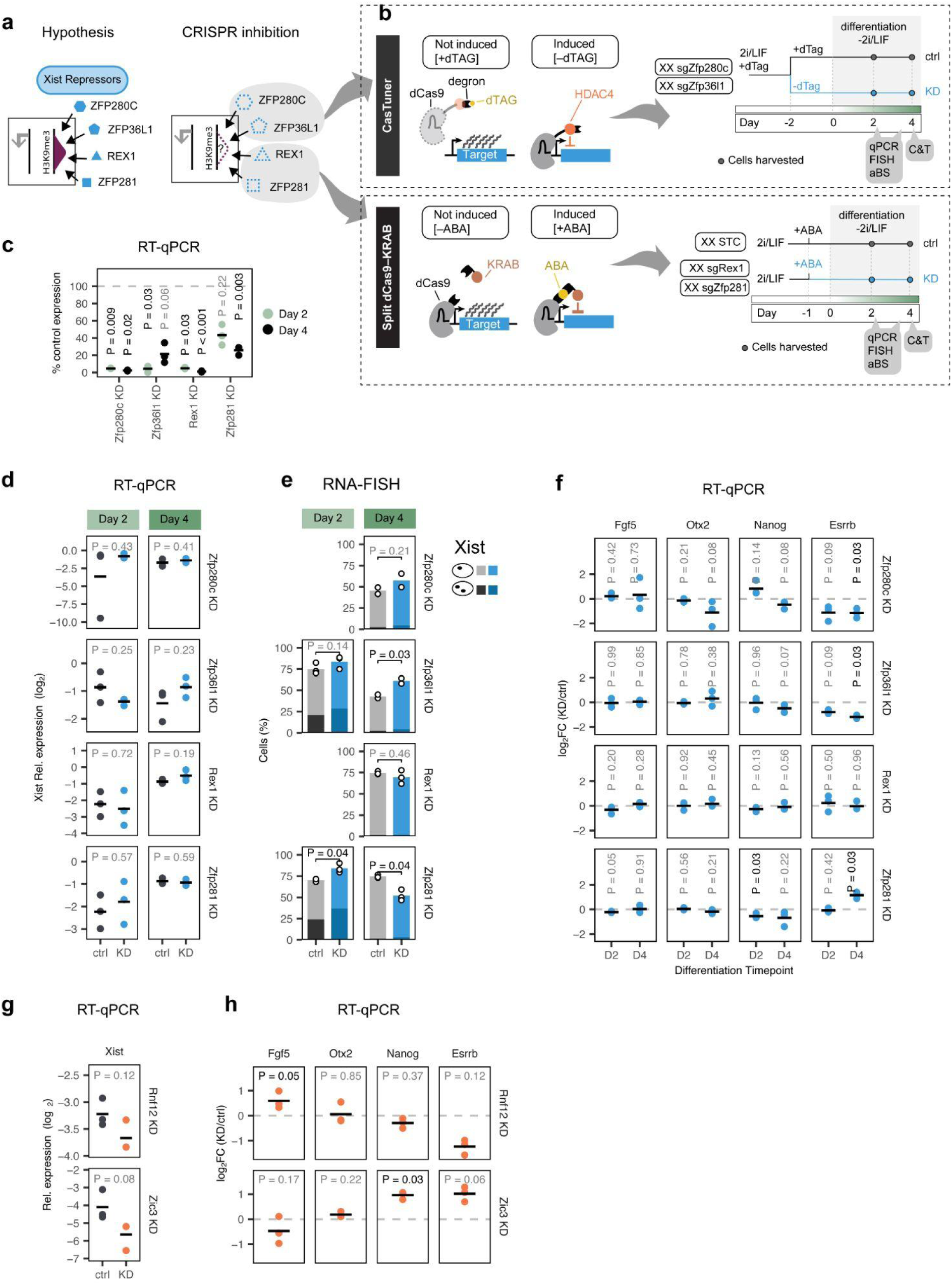
(a) Schematic representation of the hypothesis tested in b-f: Xist repressors (blue) might function by promoting H3K9me3 (purple) at the *Xist* locus. This was tested by depleting repressors by CRISPRi. (b) Overview of CRISPRi systems and experimental setup used. Top: CasTuner (dCas9-HDAC-degron) is degraded in the presence of dTAG and KD is induced by dTAG removal. Bottom: With the split dCas9-KRAB system, a KRAB repressor domain is tethered to dCas9 in the presence of ABA, and a safe-targeting control (STC) guide is used as control. aBS: amplicon-bisulfite sequencing. C&T: CUT&Tag. (c) KD efficiency measured by RT-qPCR. The black lines represent the mean of the 3 biological replicates. Two-sided unpaired T-test of relative expression of KD vs control, reported p-value black if significant (P < 0.05). (d) RT-qPCR for Xist for each of the KD cell lines (rows), at differentiation days 2 and 4. Results of a two-sided unpaired T-test are indicated. (e) RNA-FISH for Xist upon repressor KD for differentiation day 2 (left, n=3 biological replicates) and day 4 (right, biological replicates n=2 for Zfp280c ctrl/KD and Zfp36l1 ctrl/KD, n=3 for STC ctrl, Rex1 KD, Zfp281 KD). Two-sided paired T-test. 100-130 cells counted from each sample. Monoallelic cells with lighter shade, biallelic with darker shade. (f) RT-qPCR to assess effect of the KD on pluripotency (Nanog, Esrrb) and differentiation (Otx2, Fgf5) markers at differentiation day 2 and 4 (D2, D4). log_2_ foldchange of the KD relative to the control. Horizontal bars represent the mean of 3 biological replicates (dots). Two-sided unpaired T-test, p-value reported (P), black for P<0.05. (g) RT-qPCR for Xist for each of the activator KDs at differentiation day 3. The black lines represent the mean of the 3 biological replicates. Two-sided unpaired T-test, p-value reported (P), black for p-value <0.05. (h) RT-qPCR to assess effect of the Rnf12 and Zic3 KD on pluripotency (Nanog, Esrrb) and differentiation (Otx2, Fgf5). log_2_ foldchange of KD relative to control. The black lines represent the mean of the 3 biological replicates. Two-sided unpaired T-test, p-value reported (P), black for p-value <0.05.

**Supplementary Figure 6:**
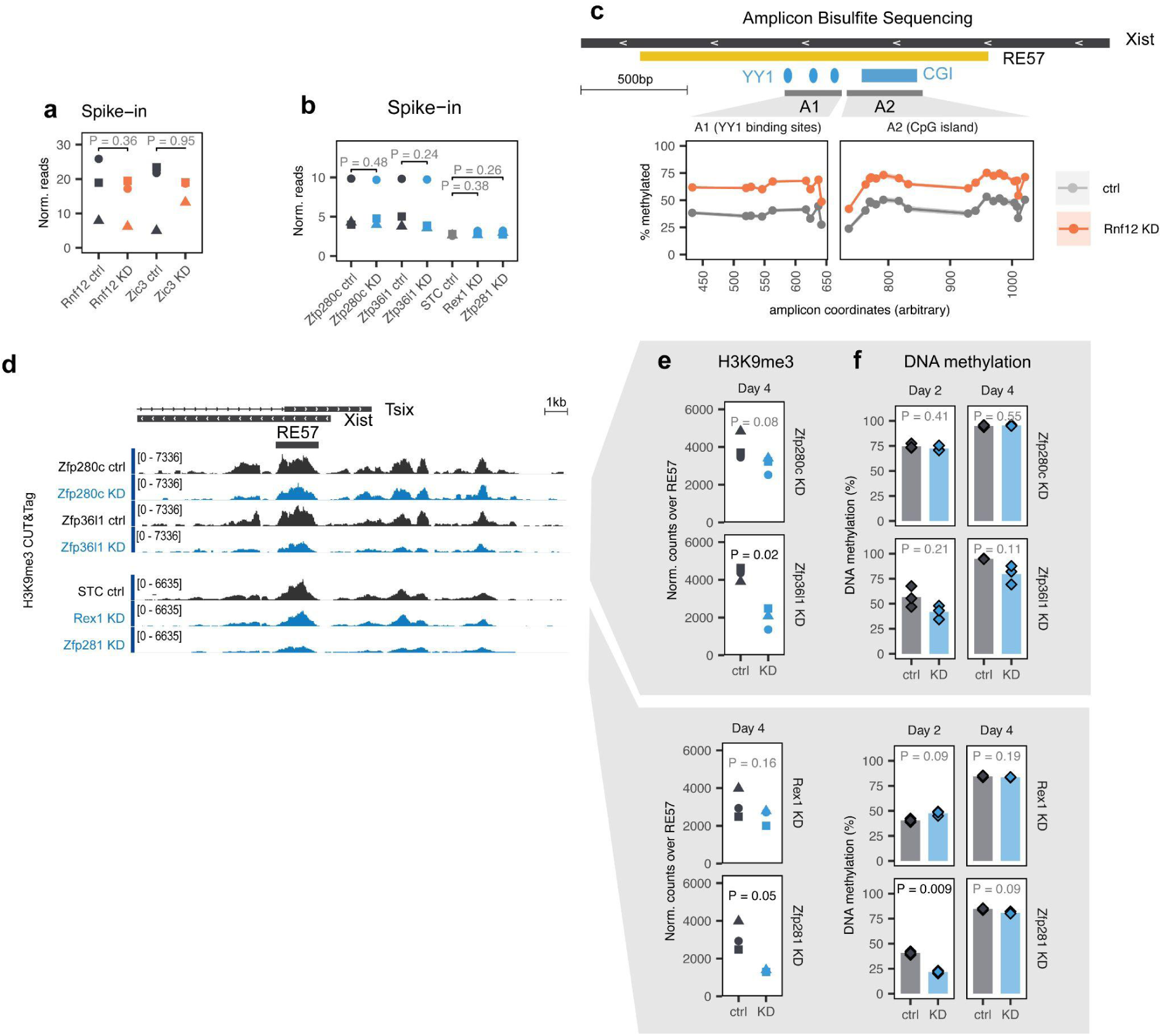
(a) Normalized spike-in reads in H3K9me3 CUT&Tag for activator KDs. Two-sided paired T-test. (b) Same as (a) for repressor KDs. (c) Amplicon bisulfite sequencing of Rnf12 KD (orange) and control (gray). Each dot is the average methylation of three biological replicates for a CpG covered by the amplicon. Ribbons indicate the confidence interval. (d) H3K9me3 CUT&Tag upon repressor KDs. Normalized tracks (average of 3 biological replicates for each condition) at chrX:103,475,052 - 103,493,155 (mm10). (e) Normalized reads covering RE57 upon repressor KDs. Two-sided paired T-test. (f) DNA methylation measured by aBS upon repressor KDs. Bars represent mean of 3 biological replicates (diamonds), p-value of two-sided paired T-test is reported.

**Supplementary Figure 7:**
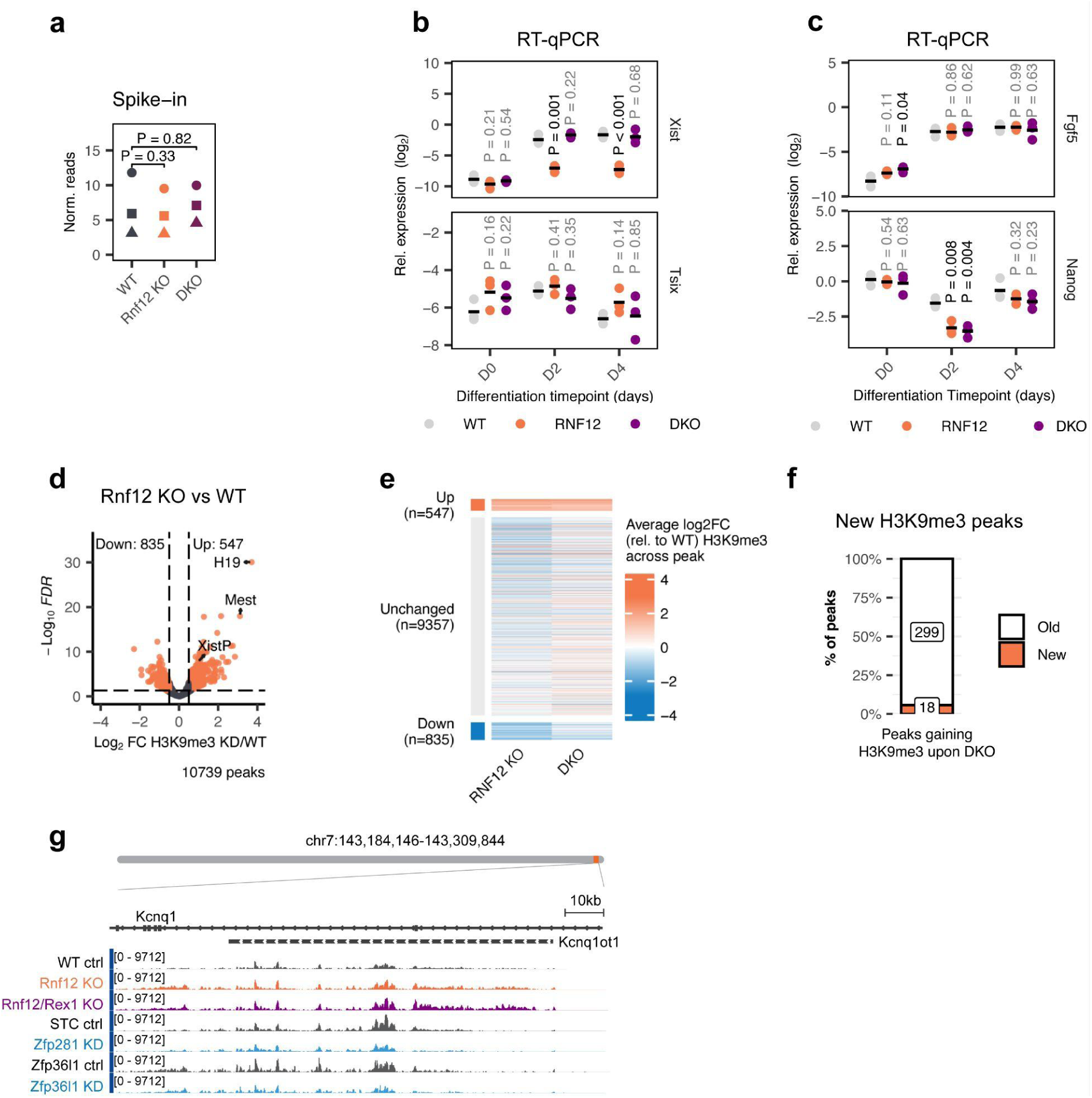
(a) Normalized spike-in reads in H3K9me3 CUT&Tag for three matched biological replicates. Two-sided paired T-test. (b) RT-qPCR for each of the KO cell lines to assess the effect of the KD on Xist and Tsix. Horizontal lines represent mean of 3 biological replicates (dots). Two-sided unpaired T-test, p-value black if p<0.05. (c) RT-qPCR to assess effect of the KOs on pluripotency and differentiation. Horizontal lines represent mean of 3 biological replicates (dots). Two-sided unpaired T-test, p-value black if p<0.05. (d) Volcano plot after differential analysis of H3K9me3 CUT&Tag data at the Rnf12 KO compared to the WT control (diffbind (DESeq2) on epic2-identified peaks). Orange dots represent peaks that had a significant change, defined as an FDR < 0.05 and an absolute log2 fold change > 0.5. *Xist* promoter region annotated (XistP). (e) Heatmap of average log_2_FC of H3K9me3 signal across peak. Peaks are grouped by their direction of change in the differential analysis: orange group has more H3K9me3 in Rnf12 KO compared to the parental cell line (WT), gray group is not changing significantly, while blue has less H3K9me3 in Rnf12 KO compared to WT. (f) Percentage of H3K9me3-gaining peaks in the DKO that did not exist in the WT (new peaks). The number of peaks in each category is indicated within the bar. (g) H3K9me3 CUT&Tag tracks at the *Kcnqot1* locus.

